# Accelerated cognitive decline in obese mouse model of Alzheimer’s disease is linked to sialic acid-driven immune deregulation

**DOI:** 10.1101/2022.02.05.479219

**Authors:** Stefano Suzzi, Tommaso Croese, Adi Ravid, Or Gold, Abbe R. Clark, Sedi Medina, Daniel Kitsberg, Miriam Adam, Katherine A. Vernon, Eva Kohnert, Inbar Shapira, Sergey Malitsky, Maxim Itkin, Sarah P. Colaiuta, Liora Cahalon, Michal Slyper, Anna Greka, Naomi Habib, Michal Schwartz

**Author notes:** Co-first authors (Stefano Suzzi, Tommaso Croese). Co-second authors (Adi Ravid, Or Gold, Abbe R. Clark). Co-last authors (Anna Greka, Naomi Habib, Michal Schwartz). Correspondence (M.Schwartz); (N.H.); (A.G.); (S.S.).

## Abstract

Systemic immunity supports healthy brain homeostasis. Accordingly, conditions causing systemic immune deregulation may accelerate onset of neurodegeneration in predisposed individuals. Here we show that, in the 5xFAD mouse model of Alzheimer’s disease (AD), high-fat diet-induced obesity accelerated cognitive decline, which was associated with immune deviations comprising increased splenic frequencies of exhausted CD4^+^ T effector memory cells and CD4^+^FOXP3^+^ regulatory T cells (Tregs). Non-targeted plasma metabolomics identified *N*-acetylneuraminic acid (NANA), the predominant sialic acid, as the major obesity-induced metabolite in 5xFAD mice, the levels of which directly correlated with Tregs abundance and inversely correlated with cognitive performance. Visceral adipose tissue macrophages were identified by sNuc-Seq as one potential source of NANA. Exposure to NANA led to immune deregulation in middle-aged wild-type mice, and *ex vivo* in human T cells. Our study identified diet-induced immune deregulation, potentially via sialic acid, as a previously unrecognized link between obesity and AD.

## Introduction

Alzheimer’s disease (AD) is the most common form of dementia, characterized by progressive brain amyloidosis and neuronal loss. For decades regarded as neuron-driven, AD pathogenesis is now recognized as being strongly affected by the state of the whole organism^1,2^. In particular, the emerging understanding that systemic immunity is required to support brain maintenance and repair^3–11^ and that immune aging increases AD risk^12–15^ suggests that repetitive or sustained systemic immune triggers would, in the long term, exhaust peripheral immune cells and reduce their ability to protect the brain. Consistently, we and others have demonstrated in mouse models of AD that both immune deficiency and systemic immune suppression^16–20^ are associated with disease severity. Based on this cumulative evidence, we envisioned that environmental conditions that are known as strong risk factors for AD might promote disease manifestation by negatively affecting immune functions. One such environmental condition is obesity, which is one of the strongest AD risk factors and one of the most frequent AD comorbidities^21–24^.

Here, we specifically focused on diet-induced obesity, and hypothesized that its effect on AD might be mediated by systemic components that may cause systemic immune deregulation. Using a transgenic mouse model of AD (5xFAD), we found that obesity induced by high-fat diet accelerated the onset of cognitive decline, which was associated with increased splenic levels of exhausted CD4^+^ T effector memory cells and of CD4^+^FOXP3^+^ regulatory T cells, and increased abundance in the blood of the metabolite *N*-acetylneuraminic acid. *In vivo* and *ex vivo* studies revealed that this metabolite is sufficient to induce immune exhaustion.

## Results

### Diet-induced obesity accelerated AD manifestations in 5xFAD mice

To test the impact of obesity on AD pathology, we generated a mouse model of obesity-AD comorbidity by using 5xFAD mice, a transgenic model of amyloidosis^25^, that were fed with high-fat diet (HFD) or control diet (CD). Wild-type (WT) mice fed with either HFD or CD were also included in the study. Since mid-life obesity increases the risk of AD^26^, we chose a long-term diet regimen starting from around 2 up to 8-9 months of age (mo) for a total of 24-28 weeks (Fig. 1a, Extended Data Fig. 1a–h). We assayed cognitive performance longitudinally using the novel object recognition (NOR) test, which evaluates short-term object identity memory, known to decline in the 5xFAD model^27,28^ and previously shown to be negatively affected by HFD in the same model^29^ (Fig. 1b). At the age of 6.5 mo, both CD and HFD-fed 5xFAD (obese-AD) mice maintained intact novelty discrimination (Extended Data Fig. 2). At 8 mo, obese-AD mice showed loss of discrimination, whereas CD-fed 5xFAD mice still performed similarly to WT controls fed with CD or HFD (Fig. 1c). Up to the time of culling, HFD did not affect the cognitive performance of WT mice at either time point (Extended Data Fig. 2; Fig. 1c).

**Fig. 1:**
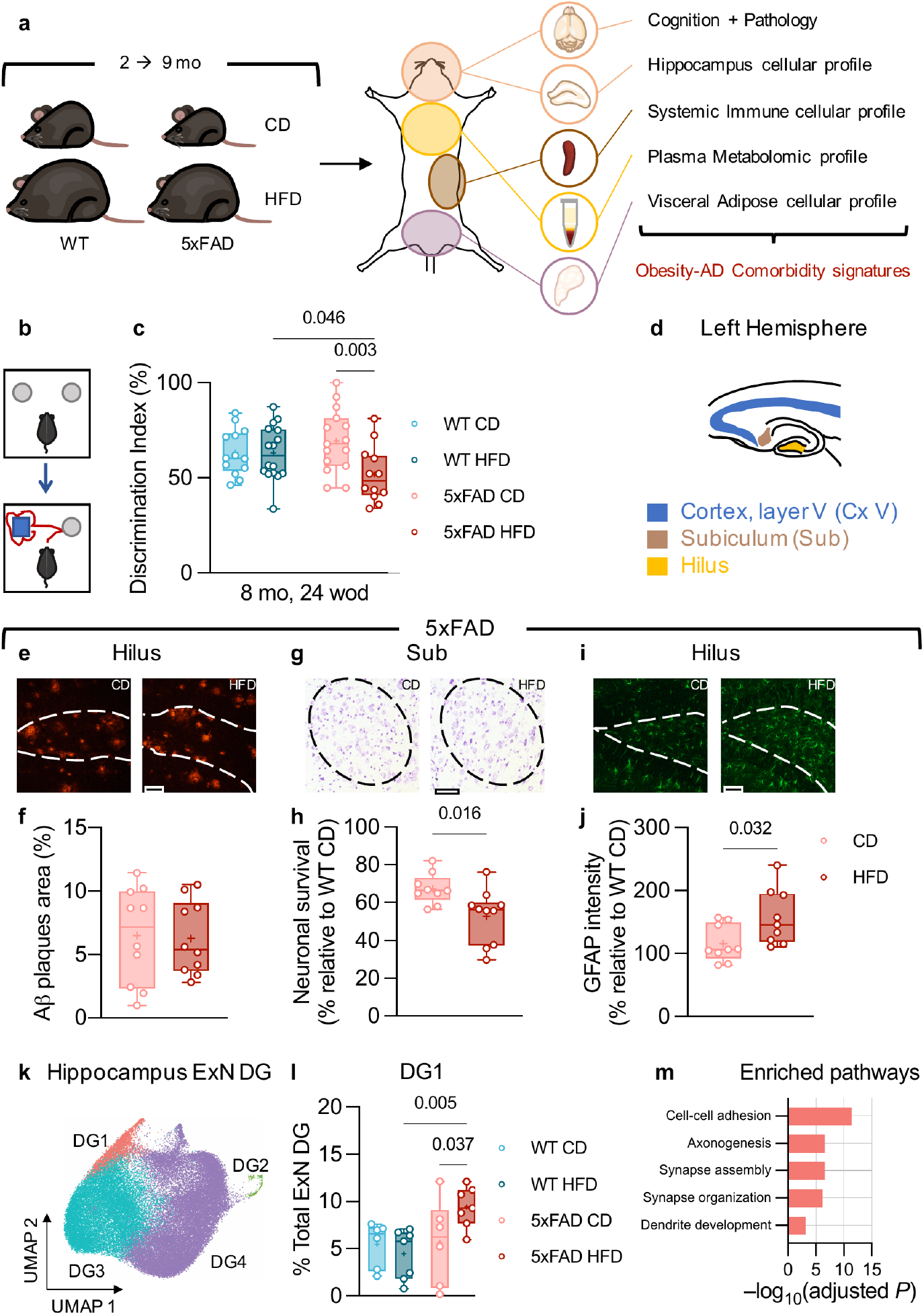
HFD leads to earlier cognitive and neuropathological manifestations in 5xFAD mice. **a**, Overview of the experimental strategy. **b**, **c**, Reduction in novelty discrimination in HFD-fed 5xFAD mice. Evaluation of cognition with the NOR task (**b**) at 8 mo, 24 weeks of diet (wod; **c**). Mice from two independent experiments, sample *n*: WT CD=13, WT HFD=16, 5xFAD CD=15, 5xFAD HFD=12. Statistical analyses: two-way ANOVA followed by Fisher’s LSD *post hoc* test. **d**, Brain regions analyzed for histopathology. **e**, **f**, Assessment of Aβ plaques load in the hippocampal hilus of 5xFAD mice. **g**, **h**, Neuronal survival in the subiculum of 5xFAD mice. **i**, **j**, Glial fibrillary acidic protein (GFAP)-immunoreactivity in the hippocampal hilus of 5xFAD mice. **e**, **g**, **i**, Representative images, left: CD, right: HFD, scale bars: 70 μm. **f**, **h**, **j**, Mice from two independent experiments, sample *n*: **f**, 5xFAD CD=10, 5xFAD HFD=10; **h**, 5xFAD CD=9, 5xFAD HFD=10; **j**, 5xFAD CD=9, 5xFAD HFD=9. **h**, Data normalized by average WT CD value (Extended Data Fig. 3b). **j**, Data normalized by average WT CD value (Extended Data Fig. 3c). Statistical analyses: two-tailed unpaired Student’s *t*-test. **k**–**m**, sNuc-Seq analysis of the hippocampus, sub-clustering analysis of the DG granule neurons (ExN DG). Mice from five independent experiments, sample *n*: WT CD=6, WT HFD=7, 5xFAD CD=6, 5xFAD HFD=7. **k**, **l**, UMAP embedding of sNuc-Seq profiles colored by cluster. **l**, Changes in frequency of DG1 cluster across experimental conditions; DG clusters 2 to 4 are shown in Extended Data Fig. 5a. Statistical analyses: two-way ANOVA followed by Fisher’s LSD *post hoc* test. **m**, Pathway analysis of the genes associated with DG1 showing enrichment of pathways related to neuronal differentiation, integration, and growth (FDR-adjusted hypergeometric test *P*-value <0.050). **c**, **f**, **h**, **j**, **l**, Box plots represent the minimum and maximum values (whiskers), the first and third quartiles (box boundaries), the median (box internal line), and the mean (cross).

Next, we assessed in the same mice if and how the obesogenic diet affected brain pathology, and specifically inspected typical AD-related hallmarks in the 5xFAD model^18–20^ (Fig. 1d). No microscopic changes were noticeable in HFD-fed WT compared to CD-fed controls (Extended Data Fig. 3a–c). In 5xFAD mice, the HFD did not affect the level of amyloid β oligomers (Aβ1-42; Extended Data Fig. 3d–f) nor amyloid β plaques (Fig. 1e, f; Extended Data Fig. 3g, h). However, compared to CD-fed 5xFAD mice, the HFD-5xFAD mice displayed increased neuronal loss (Fig. 1g, h; Extended Data Fig. 3i, j) and earlier manifestation of astrogliosis relative to CD-5xFAD mice (Fig. 1i, j). Altogether, these results indicate that the HFD accelerated the onset of disease manifestations in the 5xFAD model of AD but did not have any effect on otherwise healthy age-matched WT mice.

The earlier onset of cognitive deficits and the exacerbated brain pathology in obese-AD mice relative to CD-fed 5xFAD mice spurred us to more closely analyze the brain’s cellular landscape. To this end, we performed single-nucleus RNA-sequencing of the hippocampus (sNuc-Seq, 10x genomics, Methods; Extended Data Fig. 4a–f, 5a–g). Briefly, we found effects of both morbidities, most prominently AD-associated alterations in microglia and astrocytes in 5xFAD mice, as previously reported^30,31^, and HFD-associated decrease in distinct neuronal types in WT mice (ExN CA1 and GABA; Extended Data Fig. 4f). Interestingly, we found a cluster of cells within the dentate gyrus (DG1) that was specifically over-represented in obese-AD mice (Fig. 1k, l; Extended Data Fig. 5a). Gene set enrichment analysis of this cluster highlighted several pathways related to neuronal differentiation, integration, and growth (Fig. 1m; Extended Data Table 1), indicative of an immature-like neuronal phenotype. Yet, in contrast to AD, HFD did not have a major qualitative effect on the cellular landscape of the hippocampus that could explain the rapid progression of the disease in obese-AD mice, arguing in favor of an HFD-induced catalyzer outside the brain.

### HFD perturbed the immune system of 5xFAD mice

Obesity is a complex systemic disease which alters many processes and pathways, any of which could affect the immune system^32–34^. To test our working hypothesis that HFD-induced obesity might accelerate disease manifestation via imposing systemic immune deregulation, we analyzed freshly isolated splenocytes from mice culled after 28 weeks of diet by flow cytometry (Extended Data Fig. 6a). Strikingly, we found shrinkage of the naive CD4^+^ T-cell population (Fig. 2a) and expansion of CD4^+^ T effector memory cells (TEMs; Fig. 2b) and CD4^+^FOXP3^+^ regulatory T cells (Tregs; Fig. 2c), while TCRβ^+^CD4^−^ T cells were unaffected (Extended Data Fig. 6b–d).

**Fig. 2:**
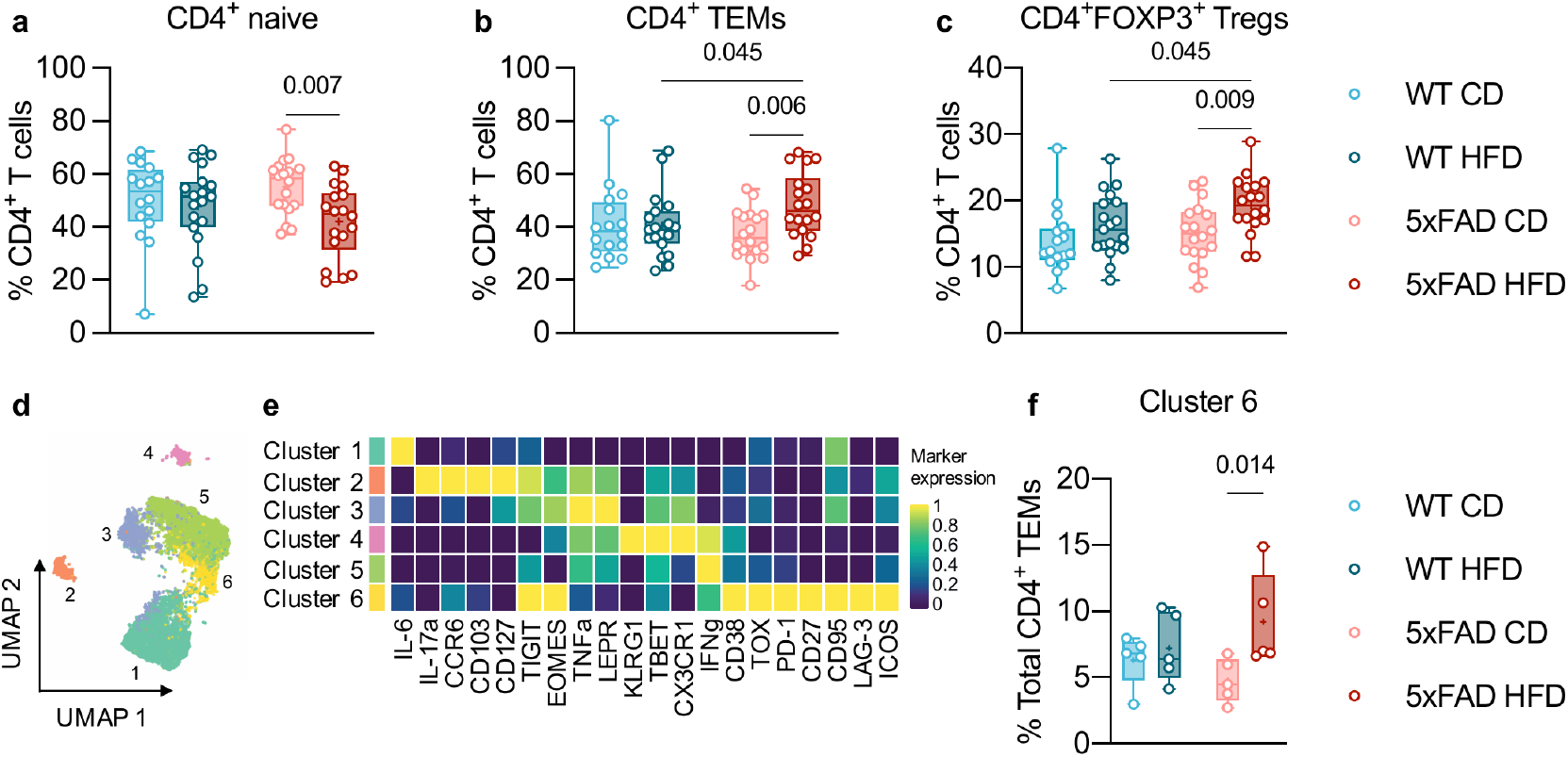
Immune-cell deviations caused by HFD in 5xFAD mice. **a**, **c**, CD4^+^ T-cell immune deviations in HFD-fed 5xFAD mice. Flow cytometric quantification of splenic frequencies of CD4^+^ naive T cells (CD44^low^CD62L^high^; **a**), CD4^+^ TEMs (CD44^high^CD62L^low^; **b**), and CD4^+^FOXP3^+^ Tregs (**c**). Mice from two independent experiments, sample *n*: WT CD=16, WT HFD=19, 5xFAD CD=18, 5xFAD HFD=18. Statistical analyses: two-way ANOVA followed by Fisher’s LSD *post hoc* test. **d**–**f**, Characterization of the CD4^+^ TEM compartment by mass cytometry. Cryopreserved splenocytes from mice evaluated for both cognition (NOR task; Fig. 1c) and systemic immune phenotype (Fig. 2a–c) were used for CyTOF analysis of the CD4^+^ TEM compartment, sample *n*: WT CD=5, WT HFD=5, 5xFAD CD=5, 5xFAD HFD=5. Cryopreserved splenocytes were restimulated with phorbol 12-myristate 13-acetate/ionomycin. **d**, UMAP embedding of CD4^+^ TEM cell clusters (2,000 cells, randomly selected from each animal). FlowSOM-based immune cell populations are overlaid as a color dimension. **e**, Mean population expression levels of markers used for UMAP visualization and FlowSOM clustering of CD4^+^ TEMs. **f**, Increased frequency of exhausted IFNg-producing TEMs in HFD-fed 5xFAD mice. Sub-clustering analysis of the CD4^+^ TEM compartment identified six clusters; Cluster 6 only is shown here, Clusters 1 to 5 are shown in Extended Data Fig. 8a. Statistical analyses: two-way ANOVA followed by Fisher’s LSD *post hoc* test. **a**–**c**, **f**, Box plots represent the minimum and maximum values (whiskers), the first and third quartiles (box boundaries), the median (box internal line), and the mean (cross).

T-cell exhaustion is one of the many age-related changes in the immune system, marked by the progressive loss of effector functions, including cytokine production^35^. Phenotypically, exhausted cells are characterized by the high and prolonged expression of inhibitory immune checkpoint receptors, and particularly the programmed cell-death protein 1 (PD-1)^36^. In addition, exhausted TEMs display a distinct transcriptional and epigenetic landscape driven by the transcription factor TOX^37^. We therefore used mass cytometry (CyTOF) to investigate the expression of exhaustion-associated markers on the CD4^+^ TEM compartment on cryopreserved mouse splenocytes (Fig. 2a– c). Sub-clustering analysis of the CD4^+^CD44^high^ TEM population (Extended Data Fig. 7a–c) identified 6 distinct clusters (Fig. 2d–f; Extended Data Fig. 8a). Interestingly, obese-AD mice displayed the highest frequencies of exhausted TEMs (Cluster 6), characterized by reduced expression of interferon gamma (IFNg) and tumor necrosis factor alpha (TNFa) and increased expression of the inhibitory immune checkpoint receptors PD-1 and LAG-3^38^, the transcription factor TOX, and other exhaustion markers, such as CD38^39^ and EOMES^40^ (Fig. 2e, f). Furthermore, analysis of the expression of the same exhaustion markers over the entire CD4^+^ TEM compartment also revealed a specific increase in obese-AD mice (Extended Data Fig. 8b). Overall, the 5xFAD mouse immune system showed higher sensitivity to the HFD-induced challenges relative to age-matched wild type mice, which resulted in increased Tregs and PD1^+^ exhausted TEMs, reminiscent of systemic immune ageing.

### Plasma *N*-acetylneuraminic acid linked cognitive loss and immune changes

We hypothesized that diet-induced metabolites might be responsible for the specific effect of HFD on the immune system of obese-AD mice. To probe this question, we carried out non-targeted polar metabolite analysis of the plasma collected from mice that were evaluated for both cognition and systemic immune phenotype. We identified a total of 229 metabolites. For each of the identified metabolites, a cell means model was built to describe the relationship between metabolite levels in the plasma and each of the genotype:diet conditions (WT:CD, WT:HFD, 5xFAD:CD, and 5xFAD:HFD; Methods). We identified 46 metabolites with significant differential levels between conditions (*omnibus* ANOVA test *P*-value <0.050, Fig. 3a). Of these, 22 were significant (*P*-value <0.050) for the 5xFAD:HFD condition, and were further tested for potential association with the cognitive score (i.e. the NOR discrimination index), on one hand, and the splenic levels of Tregs, on the other hand (Fig. 3b–d). *N*-acetylneuraminic acid (NANA), the predominant form of sialic acid in mammals^41^, was the metabolite that inversely correlated with cognitive performance (Fig. 3c) and directly correlated with increased Treg frequency (Fig. 3d). The levels of free, unbound NANA were also independently validated using a fluorometric assay on a larger number of samples and were again found to be the highest in obese-AD mice (Fig. 3e).

**Fig. 3:**
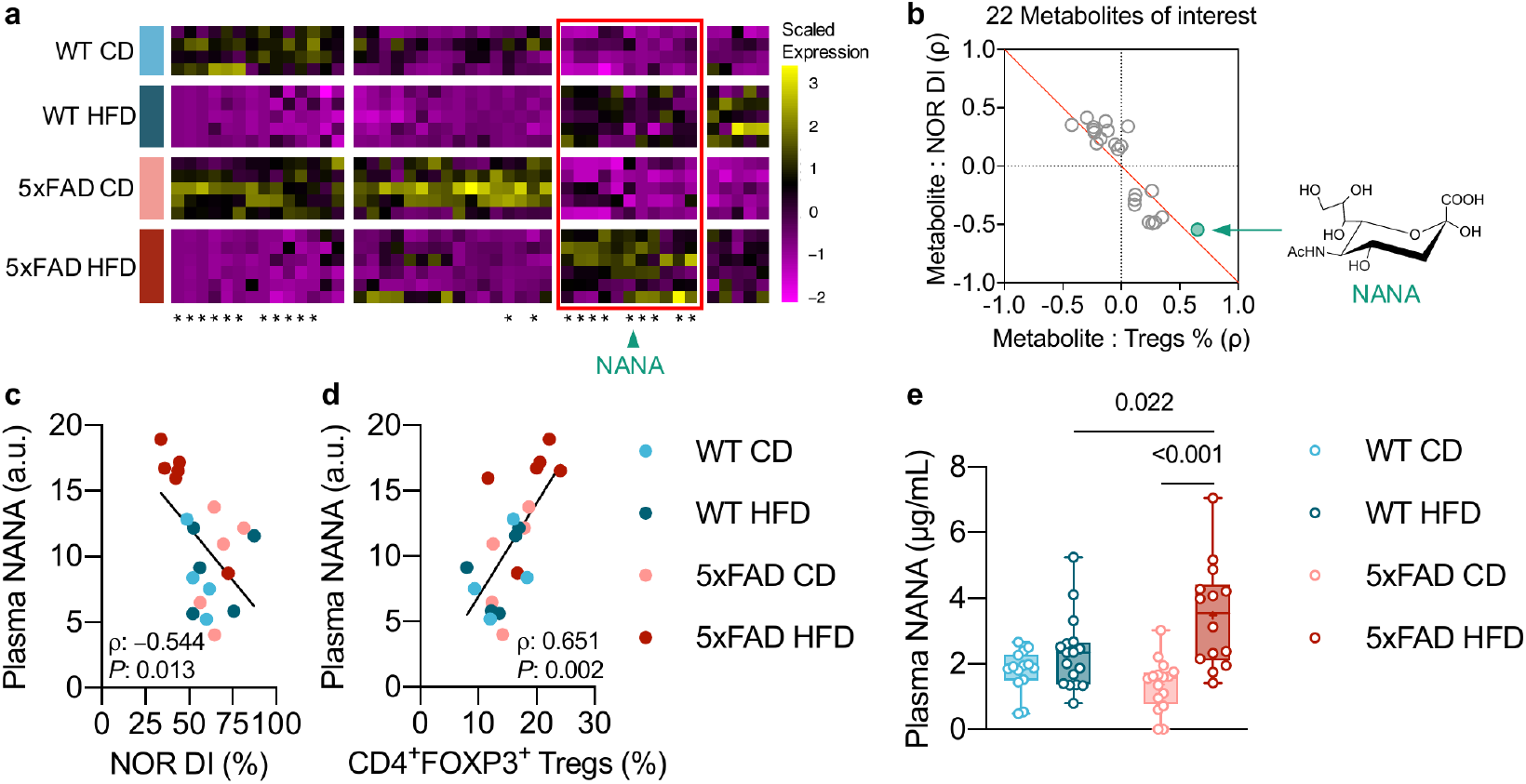
Elevated levels of circulating NANA in obese-AD mice. Non-targeted metabolomic analysis of plasma samples from mice evaluated for both cognition (NOR task; Fig. 1c) and systemic immune phenotype (Fig. 2a–c), sample *n*: WT CD=4, WT HFD=5, 5xFAD CD=5, 5xFAD HFD=6. **a**, Heatmap representation of the plasma metabolites whose linear regression models had unadjusted *omnibus* ANOVA test *P*-value <0.050 (46 out of 229 total identified metabolites). Each column represents one metabolite and each row one sample (mouse). Asterisks indicate the metabolites of interest (22 in total; Methods). The red box highlights the block of metabolites whose overall levels trended highest in HFD-fed 5xFAD mice, which include *N*- acetylneuraminic acid (NANA; green arrowhead). **b**, ρ-ρ plot (Methods). The red diagonal discriminates between metabolites associated with high NOR discrimination index (DI) and low Tregs abundance (% out of total CD4^+^ T cells; quadrant II) and metabolites associated with low NOR DI and high Tregs % (quadrant IV). The position of NANA (inset) is indicated by the green arrow. **c**, **d**, Simple linear regression (black line) and Spearman’s rank correlation (ρ coefficient, P-value) between NANA levels, as quantified after plasma metabolomic screening, and NOR discrimination index (DI; **c**) and Tregs abundance (**d**). **e**, Quantification of plasma NANA using a fluorometric assay. Mice from two independent experiments, also including the same animals described in **a**–**d**, sample *n*: WT CD=14, WT HFD=17, 5xFAD CD=16, 5xFAD HFD=14. Statistical analyses: two-way ANOVA on square root-transformed data followed by Fisher’s LSD post hoc test. Box plots represent the minimum and maximum values (whiskers), the first and third quartiles (box boundaries), the median (box internal line), and the mean (cross).

### Visceral fat is a potential source of NANA in obesity

Next, to identify a putative source of NANA, we studied the cellular landscape of the most likely target organ of the comorbidity in the periphery: the visceral adipose tissue (VAT). To this end, we analyzed the fate of the gonadal VAT by sNuc-Seq (10x genomics, Methods; Extended Data Fig. 9a–e, 10a–i). The most prominent effect among the four experimental groups was the obesity-driven expansion of macrophages, without further effect due to genotype (Extended Data Fig. 9a–e; Fig. 4a, b). In particular, sub-clustering analysis revealed a macrophage population (MAC3) that selectively expanded with HFD, characterized by the simultaneous expression of *Trem2* and several additional markers consistent with the previously described lipid-associated macrophages^42^ (LAMs; Extended Data Fig. 10a–h). In contrast, AD alone showed few discernible effects, most notably a reduction in the overall number of immune cells in CD-fed 5xFAD animals (Extended Data Fig. 9c).

**Fig. 4:**
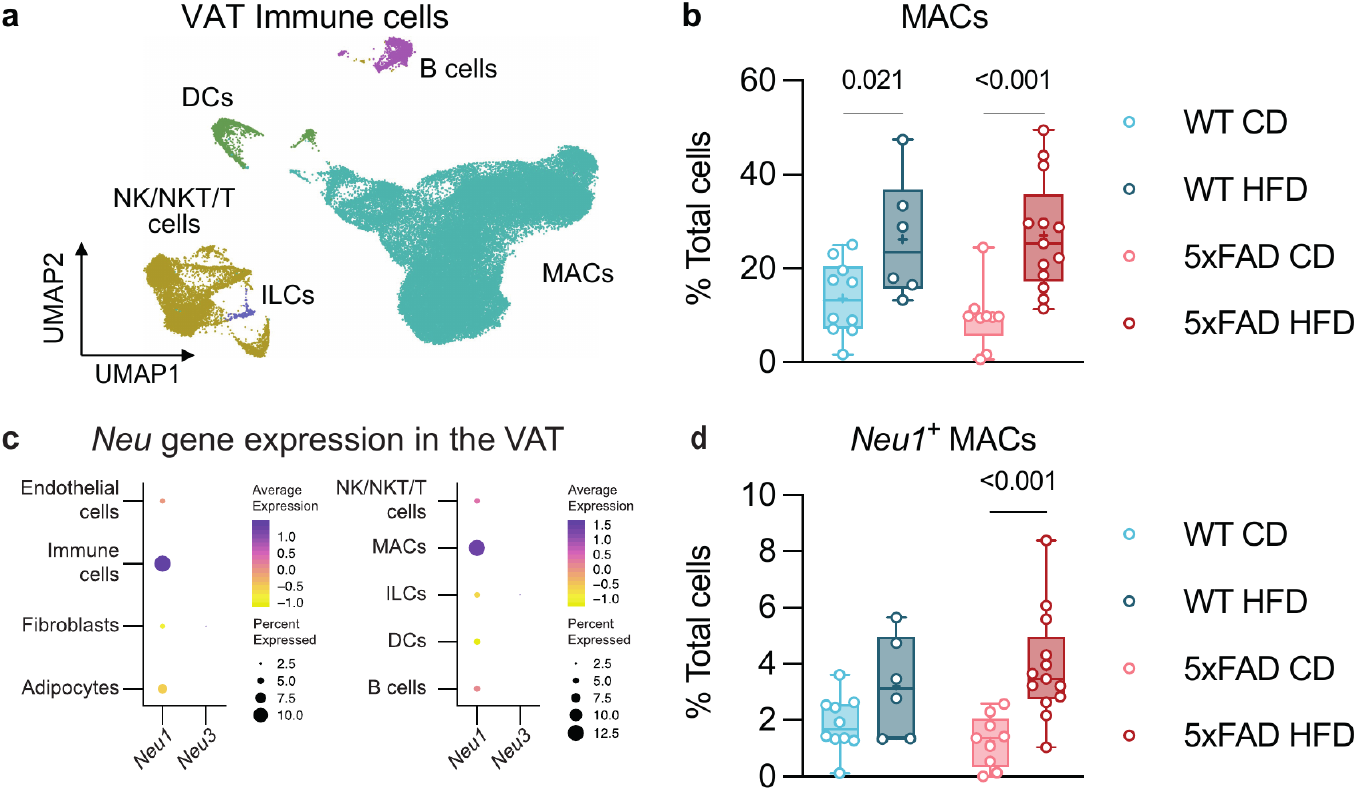
*Neu1*-expressing macrophages in the VAT are potential source of NANA. **a**, Cellular landscape of the mouse VAT immune cells across all genotype and diet conditions. UMAP embedding of single nucleus profiles (sNuc-Seq), colored after *post hoc* cell type annotation. Mice from three independent experiments, sample *n*: WT CD=10, WT HFD=6, 5xFAD CD=9, 5xFAD HFD=13. **b**, HFD increases the frequency of VAT macrophages (MACs). Changes in frequency of the other VAT immune cell types are shown in Extended Data Fig. 9e. Annotations are as in Extended Data Fig. 9d, e. **c**, Dot plots featuring the expression of sialidase *Neu* genes (color scale) and the percentage of cells expressing them (dot size) in the overall VAT (left) and VAT immune compartment (right). Of the four mammalian *Neu* genes, only *Neu1* and *Neu3* transcripts were detected. **d**, HFD increases the frequency of *Neu1*-expressing macrophages in the adipose tissue. Statistical analyses: two-way ANOVA followed by Fisher’s LSD *post hoc* test. Note: the statistics for MACs (**b**) was performed on square-root-transformed data. **b**, **d**, Box plots represent the minimum and maximum values (whiskers), the first and third quartiles (box boundaries), the median (box internal line), and the mean (cross).

Given the HFD-induced expansion of macrophages in the VAT, we sought to evaluate the expression profiles of the four neuraminidase *Neu* genes^43^, encoding for enzymes known to generate NANA. We found that *Neu1* was most abundantly expressed by immune cells, and preferentially by macrophages; in contrast, *Neu3* was hardly detectable, whereas *Neu2* and *Neu4* were below the detection level (Fig. 4c). In obese mice, the majority of *Neu1*^+^ macrophages were MAC3/LAMs (Extended Data Fig. 10i), suggesting that neuraminidase activity might be an important component in the macrophage response that was reported to be linked to disrupted adipose tissue lipid homeostasis^42^. Remarkably, *Neu1*-expressing macrophages were particularly increased in obese-AD mice (average 3.1-fold relative to CD-fed 5xFAD mice) compared to HFD-fed WT mice (average 1.7-fold relative to CD-fed WT mice; Fig. 4d). Taken together, these findings suggest that the increased proportion of *Neu1*-expressing macrophages in the VAT combined with the increase of adiposity might account, at least in part, for the specific elevation of circulating NANA levels in obese-AD mice.

### NANA drives T-cell deregulation *in vivo* and *in vitro*

To evaluate whether the increased abundance of circulating NANA could induce systemic immune deregulation, we administered NANA to regularly-fed WT mice. Given our finding that the HFD affected the immune system of 5xFAD mice, but not the WT mice, we envisioned that WT mice would not be affected by NANA unless under conditions of reduced resilience. Since chronologic aging reduces organismic resilience to stress factors^44–47^, we used both WT young adult (6.5-9 mo) and middle-aged (11-14 mo) mice and treated them with NANA, and subsequently analyzed their T-cell compartment by flow cytometry (Fig. 5a; Extended Data Fig. 11a). In young adult mice, we did not observe any effect of NANA (Fig. 5b; Extended Data Fig. 11b). In contrast, treatment of middle-aged mice with NANA caused elevation of CD4^+^ TEMs and CD4^+^PD1^+^ T cells (Fig. 5c), and marginal effect on CD8^+^ T cells (Extended Data Fig. 11c). Thus, the short-term administration of NANA in middle-aged mice was sufficient to cause immune deviations similar to those observed in obese-AD mice after months-long obesogenic diet.

**Fig. 5:**
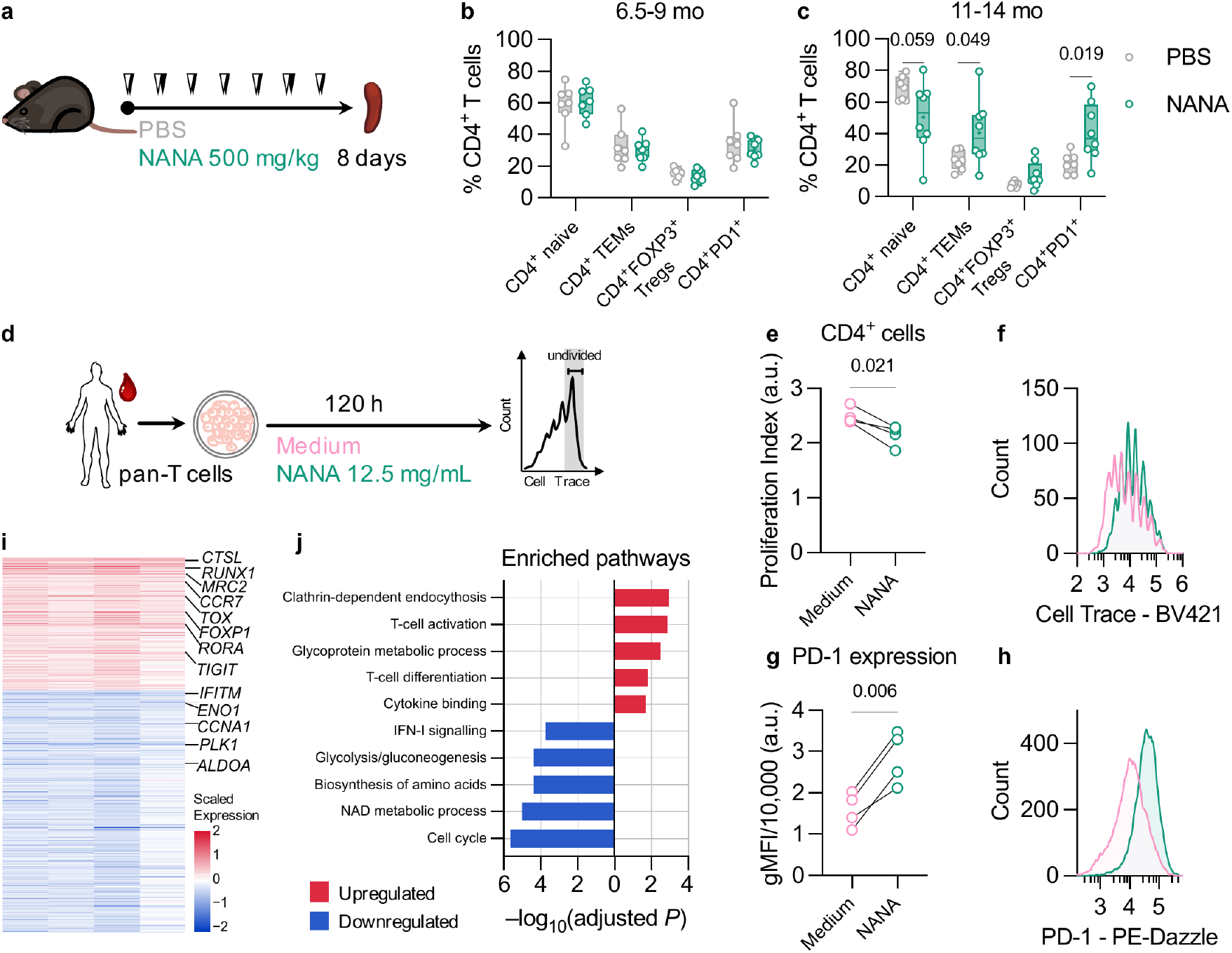
T-cell deregulation induced by NANA *in vivo* in mice, and *ex vivo* in human lymphocytes. **a**, Schematic of treatment regimen in mice. PBS control or NANA was subcutaneously injected twice a day, once in the morning (white arrowheads) and once in the evening (black arrowheads), for 7 consecutive days. **b**, **c**, Flow cytometric quantification of splenic frequencies of CD4^+^ naive T cells (CD44^low^CD62L^high^), CD4^+^ TEMs (CD44^high^CD62L^low^), CD4^+^FOXP3^+^ Tregs, and CD4^+^PD1^+^ T cells in young adult (**b**) and middle-aged mice (**c**). For both (**b**) and (**c**), data from two independent experiments, sample *n*: **b**, PBS=7, NANA=7; **c**, PBS=7, NANA=8. Statistical analyses: **b**, multiple two-tailed unpaired *t*-tests; **c**, multiple two-tailed unpaired t-tests with Welch’s correction. Box plots represent the minimum and maximum values (whiskers), the first and third quartiles (box boundaries), the median (box internal line), and the mean (cross). **d**–**h,** NANA exposure reduces human CD4^+^ T-cell proliferation and increases PD-1 expression *in vitro*. **e**, Assessment of human CD4^+^ T-cell proliferative ability. **f**, Histogram of a representative sample. **g**, Assessment of PD-1 expression (geometric mean fluorescent intensity, gMFI) in human CD4^+^ T cells. **h**, Histograms of a representative sample. **e**–**h**, Data from one experiment, sample *n*=4 human individuals (blood), each tested *ex vivo* in two conditions (NANA 12.5 mg/mL, and medium control). The presented experiment was one of three experiments with human blood samples, where different NANA concentrations were tested (Methods, *Data reporting* section). Each black line represents one individual in the two tested conditions. Statistical analyses: paired two-tailed *t*-test. **i**, **j**, Bulk RNA-Seq of the same human pan-T-cell cultures in **d** –**h**. **i**, Heatmap showing significantly (FDR-adjusted DESeq2 *P*-value <0.050) upregulated (red color scale) and downregulated genes (blue color scale). Each column represents one individual and each row represents one gene. For each individual, gene expression values are expressed as log-transformed fold-changes (NANA versus medium control). **j**, Bar plot showing significantly (FDR-adjusted hypergeometric test *P*-value <0.050) upregulated (red) and downregulated (blue) pathways.

To directly test the effect of NANA on T cells *ex vivo*, we evaluated mouse and human T-cell proliferation in the presence or absence of NANA. In both mice and humans, NANA suppressed the proliferation of CD4^+^, but not CD8^+^ T cells (Extended Data Fig. 12a–g; Fig. 5d–f; Extended Data Fig. 13a–e). In addition, NANA induced the expression of PD-1 (Fig. 5g, h) in human CD4^+^ T Cells, similar to our findings in middle-aged mice treated with NANA (Fig. 5c).

To gain mechanistic insight into the transcriptional programs linking NANA to immune deregulation, we performed bulk RNA-sequencing of the same human T-cell cultures (Extended Data Fig. 14). We found that NANA significantly upregulated 258 genes and downregulated 484 genes (DESeq2, FDR <0.050; Methods; Fig. 5i; Extended Data Table 2). Upregulated gene set enrichment analysis highlighted several pathways associated with T-cell activity, such as *clathrin-dependent endocytosis*, *T-cell activation*, and *T-cell differentiation* (FDR <0.050, Fig. 5h; Extended Data Table 3). Consistent with the results from T-cell proliferation assays, the top-downregulated pathways were related to cell proliferation (Fig. 5j; Extended Data Table 3). Moreover, we found prominent suppression of genes involved in cell metabolic and bioenergetic pathways, and specifically *NAD metabolic process*, *biosynthesis of amino acids*, and *glycolysis/gluconeogenesis* (Fig. 5j; Extended Data Table 3), thus suggesting that NANA impacts T-cell function, at least in part, via disrupted cell metabolism.

## Discussion

In the present study, we show that diet-induced obesity accelerated cognitive loss in 5xFAD mice. We further show that the comorbidity effect was functionally linked to systemic immune deregulation, which was found here to be associated with the diet-induced elevation of NANA abundance in the circulation. In independent *in vivo* and *ex vivo* experiments, NANA induced immune deregulation under seemingly low-resilience conditions, as in the 5xFAD mouse model of AD and in middle-aged WT mice, but not in young adult healthy mice.

Studies in humans and mice have shown that obesity is linked to cognitive decline^23,48^. Here, we found that 7 months of an obesogenic diet regimen did not alter cognition in young adult WT mice, but it accelerated cognitive decline in age-matched 5xFAD mice. Similarly, alterations in the systemic immune system as a result of the obesogenic diet were only observed in the 5xFAD mice, but not in WT mice. In particular, in obese-AD mice, CD4^+^ T cells underwent both quantitative and qualitative changes similar to those occurring with immune aging^35,49–51^, consistent with chronic antigenic stress and immune exhaustion^52^.

A notable finding in our study is the elevation of CD4^+^FOXP3^+^ Tregs specifically in obese-AD mice. The role of Tregs in neurodegeneration is controversial, as both protective^53–58^ and harmful functions^18,57,59–61^ have been described, depending on location and timing^18,54,62–66^. While Tregs within the brain can display anti-inflammatory activity^18,67–71^, their elevation in the periphery might be harmful, as it blocks the ability of the brain to harness bone marrow-derived cells needed to clean the brain and resolve local inflammation^18^. In our previous work in 5xFAD mice, we have shown that brain disease progression is accompanied by increased Tregs in the periphery that interfere with recruitment of disease-resolving leukocytes into the brain by curtailing IFNg signaling at the choroid plexus^18,72^. In different mouse models (including 5xFAD), treatment that transiently reduced systemic Tregs or blocked the PD-1/PD-L1 inhibitory immune checkpoint pathway resulted in modification of disease course^18–20^ and increased Tregs levels in the brain^18,71^. In line with this previous evidence, and considering that aging, including immune aging, is the primary risk factor for AD^73^, the most parsimonious interpretation of our present findings is that 5xFAD mice had reduced resilience to the obesity-induced burden on the immune system, which in turn led to the exacerbation of brain pathology.

Our analyses identified free, unbound NANA as the diet-associated metabolite that displayed the strongest association with both the cognitive decline and the immune deregulation in obese 5xFAD mice. In line with these observations, we found that a relatively aged immune system was more vulnerable to NANA. Based on this, we propose that chronic antigenic stress and systemic inflammation, which are associated with both obesity and AD, increased immune vulnerability to NANA in obese-AD mice. However, we cannot rule out the possibility that NANA had additional direct effects on the brain, for example via microglia or astrocytes, as previously reported for other metabolites^74,75^. The present findings are in line with the reported elevation of total sialic acid in the circulation in a wide range of AD comorbidities, such as aging^76^, obesity^77^, diabetes and cardiovascular disease^78^, in addition to AD itself^79^. Of note, CD4^+^ T cells of both mice and humans were more susceptible to NANA than CD8^+^ T cells, in agreement with our finding that CD4^+^ T cells were specifically perturbed in obese-AD mice.

Our data suggest that neuraminidase *Neu1*-expressing macrophages in the VAT may be a putative source of NANA in obesity. Our findings are in line with previous reports of increased NEU1 enzymatic activity in the VAT of two strains of obese mice^80^ and attenuated weight gain and VAT inflammation in diet-induced obese mice treated with a pan-neuraminidase inhibitor^81^. *Neu1* expression is required during monocyte-to-macrophage differentiation and for macrophage activation and phagocytosis^82–84^. Therefore, accumulation of *Neu1*-expressing macrophages in obese VAT may contribute to the establishment of chronic local inflammation, which may in turn lead to latent negative effects on systemic immunity.

In summary, we propose that sialic acid-driven immune deregulation may be a risk factor not only in obesity, but generally in pathological states undermining organismic resilience, and especially in compound AD-comorbid states. Our results suggest that, by accelerating systemic immune deregulation, the obesity-AD comorbidity compromises the ability of the immune system to support the brain in coping with pathology. Given the existence of numerous other environmental factors that affect the onset and severity of dementia, accelerated T-cell aging may be one common denominator. Therefore, strategies aimed at reducing immune suppression or exhaustion^18–20,85,86^ or therapies targeting diet-induced metabolites like NANA may provide salutary benefits for dementia prevention regardless of underlying etiology.

## Acknowledgements

We thank the Metabolic Profiling unit, the Targeted Metabolomics unit, Ron Rotkopf from the Bioinformatics unit, Tomer Meir Salame from the Cell Cytometry unit, Calanit Raanan from the Histology and Pathology unit at the Weizmann Institute, Michal Bronstein and her team from the Genomics Facility at the Hebrew University, and Julia Waldman and Dan Dubinsky from the Broad Institute for technical support. M.Schwartz is supported by: the Advanced European Research Council grants 232835 and 741744; the European Seventh Framework Program HEALTH-2011 (279017); the Israel Science Foundation (ISF)-research grant no. 991/16; the ISF-Legacy Heritage Bio-medical Science Partnership research grant no. 1354/15; the Thompson Foundation and Adelis Foundation. N.H. is supported by: the Israel Science Foundation (ISF) research grant no. 1709/19; the European Research Council grant 853409; the MOST-IL-China research grant no. 3-15687; the Myers Foundation. N.H. holds the Goren-Khazzam chair in neuroscience. A.G. is supported by: the National Institutes of Health (NIH) grants DK095045 and DK099465; the Cure Alzheimer’s Fund; the Chan Zuckerberg Foundation; the Carlos Slim Foundation.

## Author Contributions

S.S., T.C., A.G., N.H. and M.Schwartz designed, performed, and interpreted the experiments, and wrote the manuscript; S.S., S.Medina, T.C., and S.P.C. performed animal experiments and harvested tissues; S.S. and S.Medina performed behavioral experiments and metabolic assessment; T.C. and S.S. performed flow cytometry and mass cytometry; T.C. performed computational analysis of flow cytometry data; S.P.C. performed immunohistochemistry, imaging, and quantification; S.S. and L.C. performed Aβ1-42 ELISA; D.K. prepared RNA-Seq libraries of hippocampal samples and optimized batches; M.A. assisted in the hippocampal RNA-Seq libraries preparation, optimized nuclei isolation protocol, performed sequencing and initial pre-processing of data; A.R. and O.G. analyzed the hippocampal RNA-Seq data; S.Malitsky and M.I. performed the metabolomics experiment; S.S. analyzed the metabolomics data; K.A.V. and M.Slyper optimized the adipose nuclei isolation protocol; K.A.V., M.Slyper, and A.R.C. isolated adipose nuclei; A.R.C., E.K., and A.R. analyzed adipose RNA-Seq data; T.C. processed human blood samples and performed mouse and human lymphocyte cultures; M.A. performed the bulk RNA-Seq libraries of human lymphocyte culture and initial pre-processing of data; I.S. analyzed the bulk RNA-Seq data; S.S. performed statistics; S.S., A.R., and T.C. created figures; N.H. guided the single nuclei and bulk RNA-Seq experiments and analysis; A.G. guided the adipose RNA-Seq experiment; M.Schwartz, N.H., and A.G. conceived the study, supervised participants, and secured funding for the study.

## Methods

### Mice

All experiments detailed herein complied with the regulations formulated by the Institutional Animal Care and Use Committee (IACUC) of the Weizmann Institute of Science. Female and male mice were bred and maintained by the Animal Breeding Centre of the Weizmann Institute of Science. For comorbidity studies, heterozygous 5xFAD transgenic mice^25^ (line Tg6799, The Jackson Laboratory) on a C57/BL6-SJL background and wild-type (WT) littermates were used. Genotyping was performed by PCR analysis of ear clipping DNA, as previously described^25^. To avoid gut microbiota-related cage effects due to coprophagia^87^, 5xFAD and WT mice were housed together. For the study of the effects of NANA on the immune system *in vivo*, we used four cohorts of female and male WT mice, and specifically: one cohort of C57/BL6-SJL mice, age 6.5 mo; one cohort of C57/BL6-SJL mice, age 9 mo; one cohort C57/BL6 mice, age 11 mo; and one cohort of C57/BL6-SJL mice, age 14 mo. To avoid NANA assimilation with coprophagia, NANA-injected mice were housed separately from the PBS-injected controls. All mice were provided with regular chow (calories from proteins: 24%; calories from carbohydrates: 58%; calories from fat: 18%; 2918, Teklad), placed on a hopper integrated with the cage lid, and water *ad libitum*, and housed in cages enriched with one paper shelter. For comorbidity studies, to induce obesity, at 6-9 weeks of age, mice were switched to a high-fat diet (HFD; calories from proteins: 18%; calories from carbohydrates: 22%; calories from fat: 60%; TD.06414, Teklad), and the food pellet checked twice a week for replenishment. Non-obese controls were kept on regular chow (control diet, CD). Mice allocated for behavioral studies or NANA/PBS injections were switched to a 24 h reversed dark-light cycle (lights on from 8 pm to 8 am) at least 7 days prior to the start of the experiments, and maintained in the regimen until experimental endpoint.

### Glucose tolerance test

Prior to test, mice were fasted overnight for 16 h, and tail blood glucose measured for 0 time point (FreeStyle glucose meter and strips). Subsequently, mice were given 5 μL/g body weight of a 200 mg/mL glucose (Sigma-Aldrich)/water solution intraperitoneally, and blood glucose measured at 15, 30, 60, 90, and 120 min after glucose injection.

### Novel Object Recognition (NOR) task

A previously published protocol^88^ was modified as follows. Mice were handled daily for at least 7 days prior to the initiation of NOR test. The NOR test spanned 2 days and included 3 trials: habituation trial, day 1 (20 min session in the empty arena); familiarization trial, day 2 (10 min session in the arena with two identical objects located in opposite corners of the floor, approximately 9 cm from the walls on each side); test trial, day 2, 60-70 min after familiarization (6 min session in the arena with one of the objects replaced by a novel one). Familiar and novel objects were visually and tactually distinct. To control for potential positional preference, the location of the novel object relative to that of the familiar object was randomized. To control for potential object bias, the same familiar and novel objects were used for half of each experimental group and switched for the other half. Two identical arenas placed side by side were used simultaneously (one mouse in each arena). Each arena was a 41.5×41.5 cm grey plastic box. Mouse behavior was recorded and measured blindly using the Ethovision software (Noldus). Novel object preference was measured as discrimination index, expressed as time of interaction with the novel object relative to time of interaction with both objects (%) during the test trial. A discrimination index above 50% indicates novelty recognition, with 50% indicating no recognition. After each trial, the arenas and equipment were wiped with 10% ethanol. Female and male mice were tested on different days. To control for potential arena effect, mice tested in one arena at 6.5 mo were tested in the other arena at 8 mo. Each mouse was presented two different novel objects at 6.5 and 8 mo.

### Exogenous NANA administration

Mice were subcutaneously injected with 20 μL/g BW of a 25 mg/mL NANA (Biosynth-Carbosynth)/PBS solution (pH adjusted to 7.4) twice a day (8-9 h interval) for 7 consecutive days. Control mice were given PBS.

### Mouse blood and tissue collection and processing

In all experiments, sacrifices carried out in the morning (9 am – 1 pm). Mice were anesthetized, exsanguinated by heart puncture, and transcardially perfused with PBS. Blood was collected in tubes containing 20 μL heparin and spun at 3,000 g for 15 min at 4°C. Supernatant plasma was aliquoted and snap frozen in liquid nitrogen (LN). After dissection, the following brain regions were excised: for histology, left hemisphere, post-fixed in 4% paraformaldehyde (PFA)/PBS; for sNuc-Seq, left hippocampus, snap frozen in LN; for ELISA and biochemistry, right cortex and right hippocampus, snap frozen in LN. The spleen was mashed with the plunger of a syringe and treated with ammonium-chloride-potassium (ACK) lysis buffer (Gibco™) to remove erythrocytes. Splenocytes were filtered through a 70 μm nylon mesh and aliquoted: for flow cytometry and pan-T-cell cultures, aliquots were used fresh; for CyTOF, aliquots were resuspended in cell freezing medium (Sigma-Aldrich) and frozen in a Mr. Frosty container (Thermo Fisher) at −80°C. The left gonadal fat pad was snap frozen in LN for sNuc-Seq.

### Human samples

Blood from healthy volunteers was obtained after informed consent under the guidelines of the Institutional Review Board of the Rambam and Galilee Medical Centers. All subjects were free from acute infectious diseases and in good physical condition. Peripheral blood mononuclear cells (PBMCs) were freshly isolated from blood collected in EDTA-coated vials by layering diluted blood (1:1 in PBS) on top of an equal volume of Ficoll (GE Healthcare Life Sciences), followed by centrifugation and isolation of the buffy coat.

### Flow cytometry

Splenocytes were washed with ice-cold PBS and stained with LIVE/DEAD™ Fixable Aqua Dead (Thermo Fisher) according to manufacturer’s instructions. After fragment crystallizable (Fc) blocking (Biolegend), cells were stained for surface antigens (Extended Data Table 4). For intranuclear staining, a FOXP3 staining kit was used (Invitrogen) according to manufacturer’s instructions. To assess apoptosis of human lymphocytes following *ex vivo* exposure to NANA, the FITC Annexin V Apoptosis Detection Kit with PI (Biolegend) was used, according to manufacturer’s instructions. Flow cytometry data were acquired with CytExpert on a CytoFLEX system (Beckman Coulter) and analyzed using FlowJo software (v10; Tree Star). In each experiment, relevant negative, single-stained, and fluorescence-minus-one controls were used to identify the populations of interest.

### Lymphocyte cultures

Pan-T cells from mouse splenocytes and human PBMCs were isolated by negative selection using magnetic beads (Miltenyi) according to manufacturer’s instructions. From each mouse or human individual, isolated cell aliquots were plated in at least duplicates for each condition. Isolated cells were stained with CellTrace Violet Cell Proliferation Kit (Thermo Fisher) according to manufacturer’s instructions, and 8×10^4^ cells were plated into 96-well U-bottom and activated with anti-CD3/anti-CD28-coated Dynabeads (Thermo Fisher), as previously described^89^. Cells were then cultured for 96 h (mouse cell cultures) or 120 h (human cell cultures) in RPMI (BioSource International) supplemented with 5% fetal bovine serum, 5 mM glutamine, 25 mM Hepes (Sigma-Aldrich), and 1% antibiotics (Invitrogen), with or without NANA (pH adjusted to 7.4). Samples were acquired on CytoFLEX (Beckman Coulter) and analyzed using FlowJo. To assess proliferative ability, the Proliferation Tool of FlowJo was used to estimate the proliferation index, i.e. the total number of cell divisions divided by the number of cells that underwent division.

### Antibodies for flow and mass cytometry

Flow cytometry antibodies were purchased pre-conjugated to fluorophores (Extended Data Table 4). For mass cytometry, monoclonal anti-mouse antibodies (Extended Data Table 5) were purchased either pre-conjugated to heavy-metal isotopes (Fluidigm) or were conjugated using the MIBItag Conjugation protocol (IONpath). Because CD4 molecules are internalized during phorbol 12-myristate 13-acetate/ionomycin stimulation^90^, the anti-CD4 antibody was used in both the surface staining and the intracellular staining steps of mass cytometry.

### Mass cytometry

Short-term reactivation of cryopreserved splenocytes and subsequent mass cytometry analysis were performed as described previously^91^. In short, PBMCs were kept at −80°C for less than 2 months before thawing in a 37°C water bath. Cells were resuspended in cell culture medium supplemented with 1:10,000 benzonase (Sigma-Aldrich), and centrifuged at 300 g for 7 min at 24°C. Samples were then left overnight at 37°C before restimulation with 50 ng/mL phorbol 12-myristate 13-acetate (Sigma-Aldrich) and 500 ng/mL ionomycin (Sigma-Aldrich) in the presence of 1× brefeldin A (BD Biosciences) and 1× monensin (Biolegend) for 4 h at 37 °C. One anchor sample and one non-stimulated control sample were also included. Surface staining was performed for 30 min at 37°C. To identify dead cells, 2.5 μM cisplatin (Sigma-Aldrich) was added for 5 min on ice. To minimize inter-sample staining variability, sample handling time, and antibody consumption, cells were barcoded with Cell-ID 20-Plex (Fluidigm), and the combined surface- and Live/Dead-stained sample was fixed and permeabilized with FOXP3 staining kit (Invitrogen). After washing, the composite sample was incubated with 4% PFA overnight at 4°C. Prior to acquisition in a Helios™ II CyTOF® system, samples were washed with cell staining buffer and mass cytometry grade water. Multidimensional datasets were analyzed using Cytobank cloud-based platform, FlowJo, and R (R Core Team, 2017).

#### Acquisition and data pre-processing

Quality control and tuning of the Helios™ II CyTOF® system (Fluidigm) was performed on a daily basis and before each acquisition. Samples were acquired in two separate days, and data were normalized using five-element beads (Fluidigm) which were added to the sample immediately before every acquisition, and using an anchor sample included in each reading, as previously described^92^. For analysis, live single cells were identified based on event length, DNA (^191^Ir and ^193^Ir) and live cell (^195^Pt) channels using FlowJo. Samples were then debarcoded using Boolean gating and mass cytometry data were transformed with an inverse hyperbolic sine (arsinh) function with cofactor 5 using the R environment.

#### Algorithm-based high-dimensional analysis

After gating for CD4^+^ T cells using FlowJo, pre-processed data were considered for the analysis. All FlowSOM-based k-NN clustering was performed on the combined dataset to enable identification of small populations. For CD4^+^ TEMs, resulting nodes were meta-clustered with the indicated k-values (based on the elbow criterion) and annotated manually. FlowSOM k-NN clustering and two-dimensions UMAP projections were calculated using the CyTOF workflow package^93^.

### Plasma preparation for metabolite profiling

Plasma samples from some of the male mice included in the experiments described in Fig. 1c and in Fig. 2a–c were used. Extraction and analysis of polar metabolites were performed as previously described^94,95^, with some modifications: 100 μL of plasma were mixed with 1 mL of a pre-cooled (−20°C) homogenous methanol (MetOH):methyl-tert-butyl-ether (MTBE) 1:3 (v/v) mixture. The tubes were vortexed and then sonicated for 30 min in ice-cold sonication bath (taken for a brief vortex every 10 min). Then, UPLC-grade water (DDW):MetOH (3:1, v/v) solution (0.5 mL) containing internal following standards: ^13^C- and ^15^N-labeled amino acids standard mix (Sigma-Aldrich) was added to the tubes followed by centrifugation. The upper organic phase was transferred into 2 mL Eppendorf tube. The polar phase was re-extracted as described above, with 0.5 mL of MTBE. Both organic phases were combined and dried in speedvac and then stored at −80°C until analysis. Lower, polar phase used for polar metabolite analysis was lyophilized, dissolved in 150 μL of 1:1 DDW:MetOH (v/v), centrifuged at 20,000 g for 5 min, transferred to a new tube, and centrifuged again. For analysis, 80 μL were transferred into HPLC vials.

### LC-MS polar metabolite analysis

Metabolic profiling of polar phase was done as previously described^95^ with minor modifications described below. Briefly, analysis was performed using Acquity I class UPLC System combined with mass spectrometer Q Exactive Plus Orbitrap™ (Thermo Fisher) that was operated in a negative ionization mode. The LC separation was done using the SeQuant Zic-pHilic (150×2.1 mm) with the SeQuant guard column (20×2.1 mm; Merck). The composition of mobile phase B and A was acetonitrile, and 20 mM ammonium carbonate with 0.1% ammonium hydroxide in DDW:acetonitrile (80:20, v/v), respectively. The flow rate was kept at 200 μL/min and gradient as follows: 0-2 min 75% of B, 14 min 25% of B, 18 min 25% of B, 19 min 75% of B, for 4 min.

#### Polar metabolites data analysis

The data processing was done using TraceFinder (Thermo Fisher). The detected compounds were identified by accurate mass, retention time, isotope pattern, fragments and verified using in-house-generated mass spectra library. Peaks were quantified by calculating the area under curve (AUC) and then normalizing the AUC values by internal standards and original sample volume. A total of 229 metabolites were identified.

#### Metabolites selection for follow-up studies

For each of the identified metabolite, a linear regression model was built with metabolite levels in the plasma across all samples of all experimental conditions as the dependent variable, and the interaction between the two categorical “genotype” and “diet” variables (“genotype:diet” term) as the only predictor. To avoid collinearity, the intercept was set to 0. The overall approach is equivalent to a cell means model, which fits an individual mean for each of the predictor levels. The metabolites whose models had unadjusted *omnibus* ANOVA test *P*-value <0.050 and *P*-value for the 5xFAD:HFD condition <0.050 were considered “metabolites of interest”. The metabolites of interest were, in total, 22. For each metabolite of interest, the correlation between the metabolite’s levels and cognitive performance (NOR discrimination index) or Tregs abundance across all samples of all genotype:diet conditions was determined using the Spearman’s rank correlation. For each metabolite of interest, the coefficients of the Spearman’s rank correlations (ρ) between metabolite level and NOR discrimination index (DI), and between metabolite level and Tregs abundance, across all samples of all genotype:diet combinations, were calculated and plotted against each other (ρ-ρ plot).

### Fluorometric assays on plasma samples

Assays were performed using commercially available kits (Abcam) according to manufacturer’s instructions, but scaling reaction volumes 1:5 to allow running in 384-well flat-bottom black plates (Greiner). To measure total cholesterol, the cholesterol assay kit – HDL and LDL/VLDL (ab65390) was used. To measure free NANA, the sialic acid (NANA) assay kit (ab83375) was used, and reactions, including background controls, were run for 1 h at RT. Samples were run in duplicates. Data were acquired using an Infinite 200 PRO microplate reader (Tecan) or a Spark microplate reader (Tecan).

### Measurement of plasma leptin

The mouse leptin ELISA kit (Abcam, ab100718) was used according to manufacturer’s instructions. Samples were run in duplicates. Data were acquired using an Infinite 200 PRO microplate reader (Tecan).

### Nuclei isolation and single-nucleus RNA library preparation

#### Brain samples

Hippocampus tissue specimens were kept frozen at −80°C until processing. Samples were batched in sets of four representing all experimental and sex groups. Working on ice throughout the nuclei isolation process, the frozen hippocampus tissue was transferred into a Dounce homogenizer (Sigma-Aldrich, D8938) with 2 mL of lysis buffer. Lysis buffer used was either EZ Lysis Buffer, as we previously used^96^ (Sigma-Aldrich, NUC101), or Igepal Lysis Buffer, containing 0.1% IGEPAL^®^ CA-630 (Sigma-Aldrich, I8896), 10 mM Tris HCL pH 7.5, 146 mM NaCl, 3 mM MgCl_2_, 40 U/mL of RNAse inhibitor (NEB M0314L, as we previously used^31^). Tissue was gently homogenized while on ice 15 times with pestle A followed by 15 times with pestle B, then transferred to a 15 mL conical tube. A further 3 mL of lysis buffer was added to a final volume of 5 mL, left on ice for 5 min, and then centrifuged in a swing bucket rotor at 500 g for 5 min at 4°C. Samples were processed two at a time, the supernatant was removed, and the pellets were left on ice while processing the remaining tissues to complete a batch of 4 samples. The nuclei pellets were then resuspended in a wash buffer containing 0.02% BSA (NEB B9000S) and 40U/mL of RNAse inhibitor in PBS, and the volume adjusted to 5 mL by adding more wash buffer. The nuclei were centrifuged in a swing bucket rotor at 500 g for 5 mins at 4°C. The supernatant was removed and the pellet was gently resuspended in 500 μL of wash buffer. Nuclei were filtered through a 30 μm MACS Smartstrainer (Miltenyi, 130-098-458) and counted using the LUNA-FL™ Dual Fluorescence Cell Counter (Logos Biosystems) after staining with Acridine Orange/Propidium Iodide Stain (Logos Biosystems, F23001) to differentiate between nuclei and cell debris. Sixteen thousand (16,000) nuclei were run on the 10x Single Cell RNA-Seq Platform using the Chromium Single Cell 3’ Reagent Kits v3. Libraries were made following the manufacturer’s protocol. Briefly, single nuclei were partitioned into nanoliter-scale Gel Bead-In-Emulsion (GEMs) in the Chromium controller instrument, where cDNA shares a common 10x barcode from the bead. Amplified cDNA is measured by Qubit HS DNA assay (Thermo Fisher, Q32851) and quality assessed by High Sensitivity D5000 ScreenTape (5067-5592) with High Sensitivity D5000 Reagents (5067-5593) on the 2200 TapeStation system (Agilent). The WTA (whole transcriptome amplified) material was diluted to <8 ng/mL and processed through v3 library construction according to the manufacturer’s protocol, and resulting libraries were quantified again by Qubit and TapeStation. Libraries from 4 channels were pooled and sequenced on 1 lane of NextSeq 550 (or 8 channels sequenced on two NextSeq 550 runs) at the Center for Genomic Technologies in the Institute of Life Sciences at The Hebrew University of Jerusalem, for a target coverage of around 150 million reads per channel.

#### Visceral adipose tissue samples

Visceral adipose tissue specimens were kept frozen at −80°C until processing. Nuclei were isolated with a previously published protocol using salt-Tris (ST)-based buffers^97^. Frozen visceral adipose tissue was placed into a gentleMACS^TM^ C tube (Miltenyi, 130-096-334) with 1 mL TST buffer and homogenized using the gentleMACS^TM^ Octo dissociator (Miltenyi). The sample was removed and with a further 1 mL of TST, was transferred to a 15 mL conical tube on ice for 10 min. Next, the sample was filtered through a 40 μm Falcon^TM^ cell strainer (Thermo Fisher, 08-771-1), to which 3 mL of 1x ST buffer was also added through the filter. The resulting 5 mL sample was then centrifuged at 500 g for 5 min at 4°C in a swinging bucket centrifuge, following which the supernatant was removed and the pellet re-suspended in 1x ST buffer (volume determined by pellet size). The nuclei-containing solution was transferred to a 5 mL polystyrene tube through a 35 μm cell strainer cap (Thermo Fisher, 08-771-23), and nuclei were counted using a C-chip disposable hemocytometer (VWR, 82030-468). Eight thousand (8,000) single nuclei were loaded into each channel of the Chromium single cell 3’ chip, for V3 10x technology (10x Genomics). Single nuclei were partitioned into GEMs and incubated to generate barcoded cDNA by reverse transcription. Barcoded cDNA was next amplified by PCR prior to library construction. Libraries of paired-end constructs were generated using fragmentation, sample index and adaptor ligation, and PCR according to the manufacturer’s recommendations (10x Genomics). Libraries from four 10x channels were pooled together and sequenced on one lane of an Illumina HiSeq X (Illumina) by the Genomics Platform of the Broad Institute.

### Quality controls for sequencing and pre-processing of sNuc-Seq data

De-multiplexing of samples after Illumina sequencing was done using 10x Cellranger version 5.0.0 mkfastq to generate a Fastq file for each sample. Alignment to the mm10 transcriptome and unique molecular identifier (UMI)-collapsing were performed using the Cellranger count (version 5.0.0, mm10-2020-A_premrna transcriptome, single cell 3’ chemistry). Separate Fastq files of the same mouse sample were combined by running the CellRanger count with multiple fastqs input parameters. Since nuclear RNA includes roughly equal proportions of intronic and exonic reads, we built and aligned reads to a genome reference with pre-mRNA annotations, which account for both exons and introns.

#### Technical artifacts and ambient RNA correction

To account for technical artifacts in the data, specifically correcting gene counts shifted due to ambient RNA, we ran the CellBender^98^ (version 2) program on each sample, which removes counts due to ambient RNA molecules and random barcode swapping from (raw) UMI-based single cell or nucleus RNA-Seq count matrices, and also determines which cell barcodes are valid nuclei libraries, excluding empty droplets and low quality libraries. We used the CellBender output as input to downstream analysis.

#### Data normalization

For every nucleus, we quantified the number of genes for which at least one read was mapped, and then excluded all nuclei with fewer than 100 detected genes. For adipose tissue data, nuclei with more than 5,000 genes were also filtered out. Genes that were detected in fewer than 3 nuclei were excluded. Expression values *E_i_*_,*j*_ for gene *i* in cell *j* were calculated by dividing UMI counts for gene *i* by the sum of the UMI counts in nucleus *j*, to normalize for differences in coverage, and then multiplying by 10,000 to create TPM-like values, and finally computing log_2_(TP10K + 1) using the *NormalizeData* function from the *Seurat* package^99–102^ (version 4).

#### Doublet detection

We annotated each nucleus with a doublet score – the nucleus’ probability of being a doublet, related to the fraction of artificially generated doublet neighbors (using an in-house optimization of DoubletFinder^45^ with the following parameters: PCs = 1-45, pN = 0.25, pK = 150/(#cells), pANN=False and sct=False). This score would later be considered for the removal of doublets. We first used a high-resolution clustering (1.3 for the hippocampus and 1.5 for the adipose tissue, see description under *Dimensionality reduction and clustering*). We excluded clusters that had more than 50% of cells that had over a high doublet score (0.35 for the hippocampus and 0.4 for the adipose tissue). Second, cells from other clusters that had over a high doublet score were excluded. In the WAT: 10,625 doublets were removed and 275,336 nuclei remained in the dataset. In the hippocampus: we excluded from the analysis cells from clusters classified as endothelial cells or OPCs, since the doublet detection failed for these cell types. For OPCs, these specifically include cells differentiating to oligodendrocytes. At the end of this stage in the hippocampus dataset, 38,060 doublets were removed and 269,578 nuclei remained in the data set. The downstream analysis of sub-clustering of specific cell types included a second inspection for doublets.

### Integrating datasets

#### Identification of variable genes and scaling the data matrix

After data pre-processing, samples from the four conditions within each batch were merged into a single Seurat object. For each batch, variable genes were selected by using a variance-stabilizing transformation (using FindVariableFeatures method, Seurat). This method, first, fits a line to the relationship of log(variance) and log(mean) of each gene from the non-normalized data, using local polynomial regression (LOESS); next, it standardizes the feature values using the observed mean and expected variance (given by the fitted line). Feature variance is then calculated on the standardized values and the genes with the top values were selected as variable genes for downstream analysis (2,500 for the hippocampus and 2,000 for the adipose tissue). For sub-clustering analysis of macrophages in the adipose tissue, genes in the ambient RNA signature were removed from the variable gene list to prevent clustering based on ambient RNA expression. The data was scaled, yielding the relative expression of each variable gene by scaling and centering (using ScaleData method, Seurat).

#### Data integration

After identification of variable genes per batch, the batches (7 for the hippocampus and 2 for the adipose tissue) were integrated into a single Seurat object (using Seurat v.4 integration workflow^99,100^), based on CCA algorithm, using the methods FindIntegrationAnchors followed by IntegrateData. The integrated data matrix was then used for dimensionality reduction and clustering.

### Dimensionality reduction, clustering, and visualization

The integrated data matrix (restricted to the genes chosen as integration anchors) was then used for dimensionality reduction, visualization and clustering. Dimensionality reduction was done with principal component analysis (PCA, using RunPCA method Seurat). After PCA, significant principal components (PCs) were identified using the elbow method, plotting the distribution of standard deviation of each PC (*ElbowPlot* in Seurat). In the adipose tissue analysis: 30 PCs were used. In the hippocampus analysis: 45 PCs for analysis of all cells, 20 PCs for astrocytes, 20 for microglia, and 10 for oligodendrocytes. Within the top PC space, transcriptionally similar nuclei were clustered together using a graph-based clustering approach. First, a k-nearest neighbor (k-NN) graph is constructed based on the Euclidean distance. For any two nuclei, edge weights were refined by the shared overlap of the local neighborhoods using Jaccard similarity (FindNeighbors method Seurat, with k=60). Next, nuclei were clustered using the Louvain algorithm^103^ which iteratively grouped nuclei and located communities in the input k-NN graph (FindClusters method Seurat, with resolution 0.5). Note that for the doublet detection stage on all cell types, we first used 45 PCs with a higher resolution clustering of 1.3 on data matrices that were merged based on the batch (see *Doublet detection* section). The obtained clusters were hierarchically clustered and re-ordered (using BuildClusterTree method Seurat). For visualization, the dimensionality of the datasets was further reduced by UMAP, using the same top principal components as input to the algorithm (using the RunUMAP method Seurat). Note that the distribution of samples within each cluster was examined to eliminate that clusters were driven by batch or other technical effects. Clusters with low-quality cells (low number of genes detected, and missing or low-key cell-type marker genes and house-keeping genes such as MALAT1), doublet clusters expressing markers of multiple cell types, and neuronal clusters from neighboring region of the hippocampal subiculum that appeared in an uneven form across samples, were removed from the analysis. Data visualization using UMAP showed that the clusters displayed a mixture of nuclei from all technical and biological replicates, with a variable number of genes, meaning the clustering was not driven by a technical effect.

#### Sub-clustering analysis of cells types

Specific cell types (i.e. microglia, astrocytes, oligodendrocytes, DG neurons, macrophages in the adipose tissue) were subsetted from the main dataset for a high-resolution analysis. For each such subset another cycle of clean-up was performed, removing doublet clusters based on different thresholds. Cells were clustered in high-resolution and clusters were then annotated and merged based on marker expression.

### Identification of clusters’ cell types

Identification of cell types was done in the hippocampus using an inhouse modification of a logistic regression model (linear_model.LogisticRegression from Python’s sklearn package). The modifications included calculating the classification probability of each cell, and eventually associating the cell with the original cluster, and classifying the cluster as the overall highest scoring cell type. The classifier was trained on 80% of the nuclei (∼50,000 nuclei) of our previously published and annotated sNuq-Seq data of the mouse hippocampus^31^, and tested on the remaining 20% of the nuclei. Cell types were annotated according to the classification and further validated using known marker gene and as previously published^31^. In the adipose tissue, identification of cell types was done based on known marker genes and the Bioconductor package SingleR^104^.

### Cell fraction estimations and statistics

The fraction of different cell populations (i.e. clusters) was separately computed, for each sample across all clusters, as the fraction of nuclei in each cluster out of the total number of nuclei, by a parameter of interest (e.g. diet, sex, and more). Correlations between these fractions of interest were calculated using Spearman’s rank correlation coefficient (with cor function from R’s base package, method=”spearman”). To assess whether there was a significant change between experimental groups across conditions (diet, genotype), two-way ANOVA followed by Fisher’s Least Significant Difference (LSD) *post hoc* test was used; parametric assumptions and, where appropriate, data transformations were performed as described in the *Statistical analyses* section.

### Marker genes and differential expression analysis

Marker genes for glial and dentate gyrus (DG) granule neurons sub-clusters were found using the MAST test^105^ (using the batch as a latent variable for glial cells in the brain to account for batch specific effects), which was run using the FindMarkers function in Seurat v.4 (using the assay set to *RNA* to use the normalized UMI counts values per gene). To find diet-dependent differentially expressed signatures, we used the same scheme as above using the MAST algorithms and the batch as the latent variable, within each glial cell type, comparing all cells of HFD-fed and CD-fed 5xFAD mice; similarly, WT mice were compared between diet groups. *P*-values were adjusted for multiple hypothesis testing using the Benjamini-Hochberg’s correction (FDR). Adjusted *P*-value threshold of 0.050 was used to report significant changes and a fold change threshold of 0.25.

### Bulk RNA-Seq library preparation and analysis

Cultured human T cells after treatment were flash frozen in Buffer RLT (Qiagen, 79216) and kept in −80°C until processing. Total RNA was extracted with the NucleoSpin RNA kit (Macherey-Nagel, 740955). One microgram (1 μg) total RNA was used as an input for mRNA isolation (NEB E7490S), followed by library preparation using NEBNext® Ultra™ II Directional RNA Library Prep Kit for Illumina (NEB E7760). Quantification of the libraries was done by Qubit and TapeStation. Paired-end sequencing of the libraries was performed on Nextseq 550. De-multiplexing of samples was done with Illumina’s bcl2fastq software. The fastq files were next aligned to the human genome (hg38) using STAR^106^ and the transcriptome alignment and gene counts were obtained with HTseq^107^. Aligned gene counts were normalized to fragments per kilobase of transcript per million mapped reads (log10(FPKM)) using the fpkm function in DESeq2 R package^108^ (1.32.0). Genes with low counts (<10) were filtered before performing statistical analysis. Differential expression analysis of bulk RNA-Seq data between NANA-treated samples and control was performed using the DESeq2 R package (*P*-value adjusted to multiple hypothesis testing ≤0.050), while accounting for inter-sample differences. Significantly upregulated and downregulated genes (defined as: log-fold change >0, adjusted *P*-value <0.050; and log-fold change <0, adjusted *P*-value <0.050, respectively) were determined. Up- and downregulated gene lists were separately functionally annotated with gene sets defined by the KEGG and Gene Ontology databases (org.Hs.eg.db, version 3.5.0), using enrichGO and enrichKEGG functions in R package *clusterPrifiler* (3.6.0), using the hypergeometric *P*-value and FDR correction for multiple hypothesis (with a threshold of FDR <0.050).

### Histology and immunohistochemistry

Paraffin-embedded tissue was sectioned with thickness 6 μm. One slide per animal was used for staining, each containing 5 equally spaced sections. Cresyl violet staining was performed to visualize neurons^20^. Antibody staining was performed as previously described^109^, except that all primary antibodies were incubated overnight at RT, followed by another overnight at 4°C. For Aβ staining, the Mouse On Mouse detection kit (Vector labs) was used according to manufacturer’s instructions. The following primary antibodies were used: mouse anti-human Aβ (1:150; Covance); chicken anti-GFAP (1:150; Abcam). Cy2/Cy3-conjugated anti-mouse/chicken secondary antibodies (1:150; Jackson Immunoresearch) were used. For counterstaining, 4’,6-diamidino-2-phenylindole (1:10,000; Biolegend) was used.

### Microscopic imaging and analysis

Images were acquired using a fluorescence microscope (E800, Nikon) equipped with a digital camera (DXM 1200F, Nikon), and with a ×20 NA 0.50 objective lens (Plan Fluor, Nikon). Quantitative analyses were performed by an experimenter blind to the identity of the animals, and using either the Image-Pro Plus software (Media Cybernetics) or ImageJ (NIH). Neuronal survival on cresyl violet-stained sections and Aβ plaque quantification were performed as previously described^20^. GFAP intensity was measured using the ImageJ software by applying a segmentation algorithm to mask stained areas (Otsu’s method) and subsequently measuring average integrated density over 3-5 sections per animal. For each animal, stained sections’ quantified values were averaged. Representative images were optimized using ImageJ.

### Human Aβ1-42 ELISA

Samples were homogenized in Tris Buffered Saline EDTA (TBSE) solution (50 mM Tris, 150 mM NaCl, and 2 mM EDTA, pH 7.4) with the addition of 1% Protease Inhibitor Cocktail (Sigma-Aldrich) using a microtube homogenizer with plastic pestles (for hippocampus; 1 mL/100 mg tissue) or a glass homogenizer (for cortex; 1 mL/200 mg tissue). The homogenates were then centrifuged for 40 min at 350,000 g in 500 μL polycarbonate centrifuge tubes (Beckman Coulter) at 4°C in an Optima MAX-XP Ultracentrifuge with a TLA 120.1 rotor (Beckman Coulter). The supernatant (TBSE-soluble fraction) was collected, aliquoted, and stored at −80°C until use. BCA assay (Pierce BCA Protein Assay Kit) was performed to determine total protein amount for normalization. To quantify Aβ1-42 peptides, the human Aβ42 Ultrasensitive ELISA Kit (Invitrogen) was used according to manufacturer’s instructions. Data were acquired using a Spark microplate reader (Tecan).

### Experimental design

No statistical method was used to predetermine sample sizes, which were chosen with adequate statistical power based on the literature and past experience. The specific sample sizes and tests used to analyze each set of experiments are indicated in the Figure legends. Animals were randomly allocated to experimental groups according to age, sex, and genotype. Sample selection for subsequent analyses such as CyTOF, sNuc-Seq, and metabolome profiling was based on cognitive, metabolic, flow-cytometric, and/or histological analyses. Investigators were blind to animal identity during experiments and outcome assessment, except during behavior experiments, where diet groups, but not genotypes, were obvious. No animal was excluded from analyses, except those removed before experimental endpoint according to IACUC guidelines, or because of technical reasons detailed as follows: for microscopic image analysis, poorly stained or overstained sections or slides were not included; for flow cytometry and CyTOF, samples with not enough cells to proceed with the analysis or samples in which the staining did not work were not included. For the analysis of the effects of NANA on the immune system *in vivo*, for one of the two cohorts of middle-aged mice, one PBS-injected animal was not included due to its statistics across all immune cell populations.

### Data reporting

For comorbidity studies, the data herein presented originated from five independent cohorts. Animals used for the sNuc-Seq of the hippocampus were selected from all five cohorts; animals used for the sNuc-Seq of the VAT were selected from the first three cohorts; animals used for all other analyses came from the last two cohorts. When animals from different cohorts were merged, the number of cohorts considered is indicated in the Figure legends. For *in vivo* studies with NANA, four independent experiments were conducted: two with young adult mice (6.5-9 mo), and two with elderly (11-14 mo) mice. For *ex vivo* studies with mouse lymphocytes, two independent experiments were conducted: one simultaneously testing two doses of NANA, 1 and 5 mg/mL (Extended Data Fig. 12c); and one testing 1 mg/mL of NANA. For *ex vivo* studies with human lymphocytes, we performed one pilot study to determine the optimal dose of NANA. We observed that 25 mg/mL of NANA, but not 10 mg/mL, had a suppressive effect on CD4^+^ T-cell proliferation (not shown). Since during the second experiment, at the concentration of 25 mg/mL, we observed a strong effect also on CD8^+^ T cells (not shown) but the *in vivo* (HFD-fed 5xFAD mice; NANA-injected mice) and the *in vitro* (NANA-treated pan-T-cell cultures) mouse data suggested a specific impact on CD4^+^ T cells, with only a minor effect on CD8^+^ T cell, we conducted the third experiment with the lower 12.5 mg/mL dose (Fig. 5d–h), and we further used this experimental condition for bulk RNA-Seq (Fig. 5i, j).

### Statistical analyses

Normality of data distribution was evaluated using Anderson-Darling’s, D’Agostino-Pearson’s (“*omnibus* K2”), Shapiro-Wilk’s, and Kolmogorov-Smirnov’s tests, and via visual assessment of quantile-quantile plots. In cases of strong deviation from normality and better fit with lognormal distribution, log-transformed data were used, and so indicated in Figure legends; if the whole dataset contained zero values, log(1+x) transformation was used. Homogeneity of variance was tested using *F*-test, for two groups, and Spearman’s test for heteroscedasticity, followed by visual assessment of the homoscedasticity plot, for more than two groups. Data were analyzed using two-tailed Student’s *t*-test to compare between two groups; Welch’s correction was applied for heteroscedastic groups. For lymphocyte cultures, paired *t*-test or one-way within-subjects ANOVA followed by Fisher’s LSD *post hoc* test were used. For comorbidity studies, in presence of two categorical independent variables (“genotype”, two levels: “WT” and “5xFAD”; “diet”, two levels: “CD” and “HFD”), two-way ANOVA followed by Fisher’s LSD *post hoc* test was used; square-root transformation was applied to heteroscedastic data. For weight gain and glucose tolerance test, two-way within-subjects ANOVA followed by Fisher’s LSD *post hoc* test was used. The specific data transformations and tests used are reported in the Figure legends. For null hypothesis testing, the test statistic with confidence intervals, degrees of freedom, and *P*-value(s) are reported in the GraphPad Prism “Extended Data Table 6” pzfx file. For *t*-tests and *post hoc* tests, *P*-values <0.060 are reported in the graphs, rounded to three decimal digits; *P*-value <0.050 was considered significant. Statistical analyses were carried out using GraphPad Prism version 9.0, R, and Microsoft Excel. Graphs were generated with GraphPad Prism version 9.0 and R.

### Materials availability

This study did not generate new unique reagents.

### Data availability

Raw and transformed data, statistical analyses including normality testing, null hypothesis testing, simple linear regressions and correlations, and the graphs utilized for main Figure and Extended Data Figure panels are reported in Extended Data Table 6 (file generated with GraphPad Prism). The list of identified metabolites, raw and processed sequencing and mass cytometry data will be available at the time of publication.

### Code availability

The codes used for computational analyses will be available upon acceptance of the manuscript.

## Extended Data

**Extended Data Fig. 1:**
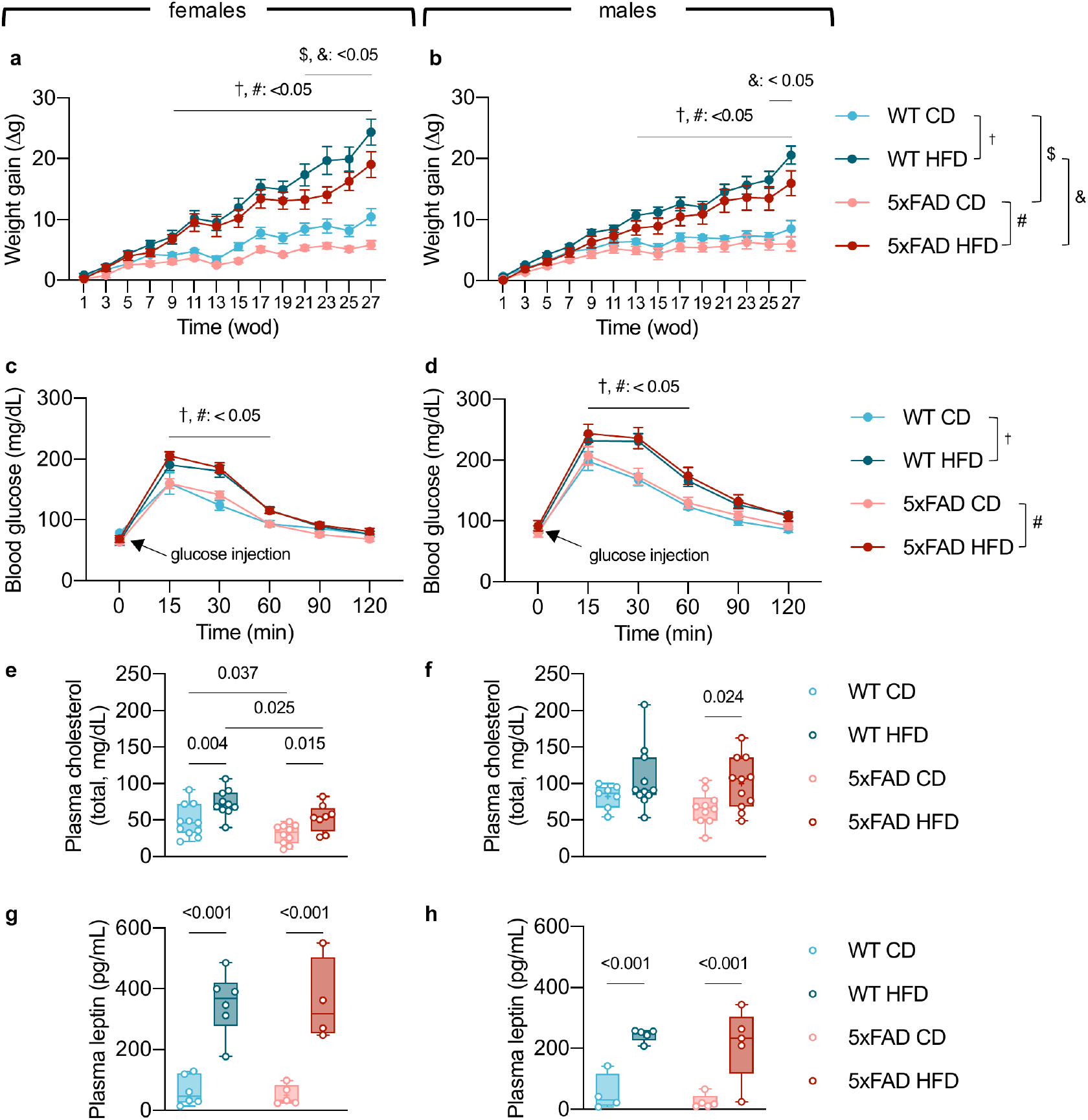
Changes in metabolism induced by HFD. **a**, **b**, Weight gain over time (weeks of diet, wod), means ± s.e.m. **c**, **d**, Glucose tolerance test, means ± s.e.m. **a**–**d**, Contrasts: †, WT CD vs. WT HFD; #, 5xFAD CD vs. 5xFAD HFD; $, WT CD vs. 5xFAD CD; &, WT HFD vs. 5xFAD HFD. **e**, **f**, Total plasma cholesterol levels. **g**, **h**, Total plasma leptin levels. **a**, **c**, **e**, **g**, Data from female mice. **b**, **d**, **f**, **h**, Data from male mice. Mice from two independent experiments, sample *n*: **a**, WT CD=9, WT HFD=10, 5xFAD CD=9, 5xFAD HFD=8; **b**, WT CD=7, WT HFD=12, 5xFAD CD=10, 5xFAD HFD=11; **c**, WT CD=11, WT HFD=10, 5xFAD CD=10, 5xFAD HFD=9; **d**, WT CD=8, WT HFD=12, 5xFAD CD=10, 5xFAD HFD=13; **e**, WT CD=11, WT HFD=10, 5xFAD CD=10, 5xFAD HFD=8; **f**, WT CD=7, WT HFD=11, 5xFAD CD=10, 5xFAD HFD=11; **g**, WT CD=6, WT HFD=6, 5xFAD CD=5, 5xFAD HFD=4; **h**, WT CD=4, WT HFD=5, 5xFAD CD=5, 5xFAD HFD=5. Some of these animals were used for the analyses described in Fig. 1–3 and Extended Data Fig. 2–8. Statistical analyses: **a**–**d**, two-way within-subjects ANOVA followed by Fisher’s LSD *post hoc* test; **e**–**h**, two-way ANOVA followed by Fisher’s LSD *post hoc* test. **e**–**h**, Box plots represent the minimum and maximum values (whiskers), the first and third quartiles (box boundaries), the median (box internal line), and the mean (cross).

**Extended Data Fig. 2:**
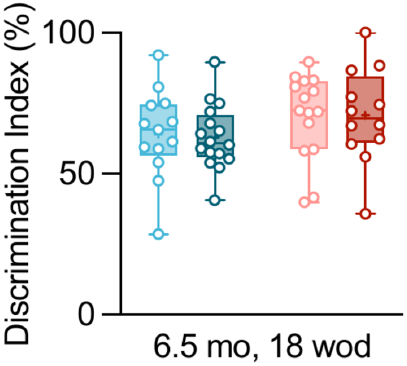
AD or HFD do not affect novelty discrimination at 6.5 mo after 18 weeks of diet (wod). Mice are the same described in Fig. 1b, sample *n*: WT CD=13, WT HFD=16, 5xFAD CD=15, 5xFAD HFD=12. Statistical analyses: two-way ANOVA followed by Fisher’s LSD *post hoc* test. Box plot represents the minimum and maximum values (whiskers), the first and third quartiles (box boundaries), the median (box internal line), and the mean (cross).

**Extended Data Fig. 3:**
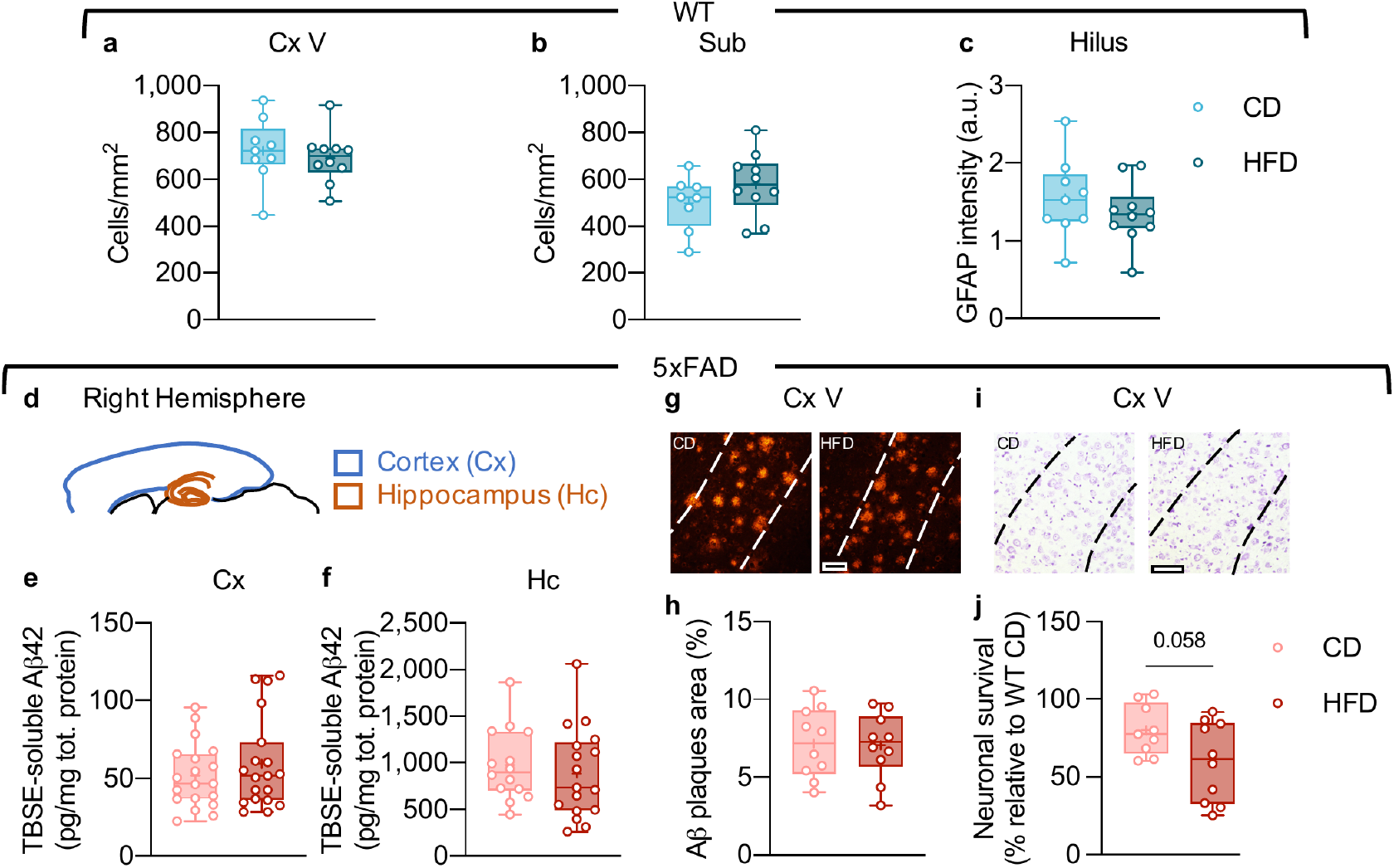
Effects of HFD on brain pathology. HFD does not affect neuronal density nor induces astrogliosis in WT mice after 28 weeks of diet. Assessment of neuronal density in the cortical layer V (**a**) and subiculum (**b**) and expression of the astrocyte marker glial fibrillary acidic protein (GFAP) intensity in the hippocampal hilus (**c**) of WT mice. Mice from two independent experiments, sample *n*: **a**, WT CD=9, WT HFD=10; **b**, WT CD=8, WT HFD=10; **c**, WT CD=9, WT HFD=10. **d**–**h,** HFD does not affect Aβ load in 5xFAD mice. **d**–**f**, Determination of Tris Buffered Saline EDTA (TBSE)-soluble Aβ1-42 oligomers in the cortex (**d**, **e**) and hippocampus (**d**, **f**) of 5xFAD mice. **g**, **h**, Assessment of Aβ plaques load. **i**, **j**, HFD does not significantly affect neuronal survival in the cortex of 5xFAD mice. **g**, **i**, Representative images, left: CD, right: HFD, scale bars: 70 μm. **e**, **f**, **h**, **j**, Mice from two independent experiments, sample *n*: **e**, 5xFAD CD=19, 5xFAD HFD=19; **f**, 5xFAD CD=14, 5xFAD HFD=17; **h**, same animals as in Fig. 1f, sample *n*: 5xFAD CD=10, 5xFAD HFD=10; **j**, same animals as in Fig. 1h, sample *n*: 5xFAD CD=9, 5xFAD HFD=10. **j**, Data normalized by average WT CD value (**a**). **a**–**c**, **e**, **f**, **h**, **j**, Statistical analyses: two-tailed unpaired Student’s *t*-test. Box plots represent the minimum and maximum values (whiskers), the first and third quartiles (box boundaries), the median (box internal line), and the mean (cross).

**Extended Data Fig. 4.**
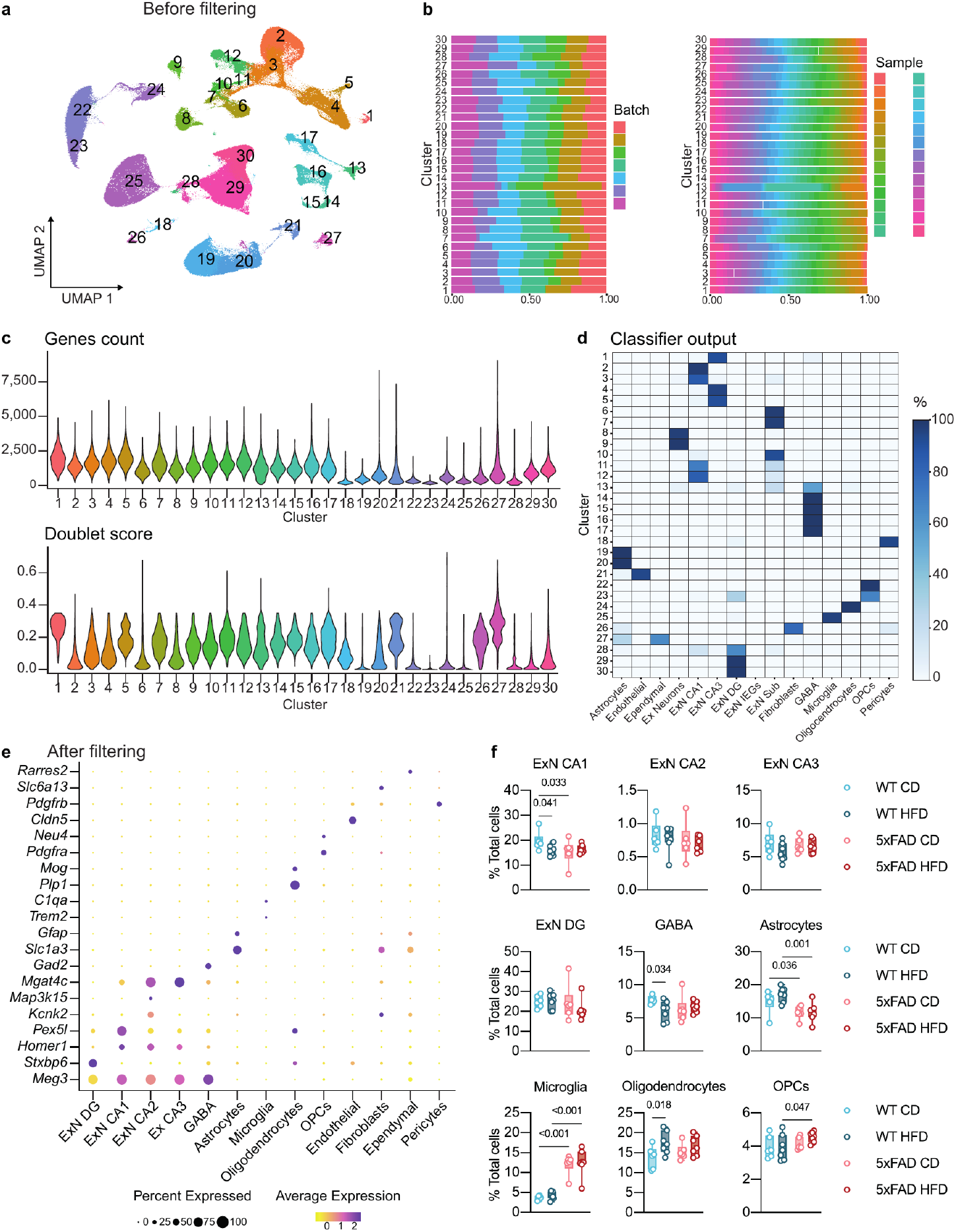
sNuc-Seq of the mouse hippocampus. Mice are the same described in Fig. 1k–m, sample *n*: WT CD=6, WT HFD=7, 5xFAD CD=6, 5xFAD HFD=7. **a**, Pre-filtered cell profiles of the mouse hippocampus across all experimental groups. UMAP embedding of single nucleus profiles colored by clusters featuring doublet cells and clusters that were excluded from the analysis. **b**, Similar distribution of cells per cluster across samples and batches. Cluster annotations are as in **a**. The percentage of cells per cluster, from the different batches (left) and from each sample (right), are colored by batch or sample. **c**, Violin plots showing the distribution of number of genes (top) and doublet score (bottom) detected in each cluster (Methods, *Quality controls for sequencing and pre-processing of sNuc-Seq data* section). Cluster annotations are as in **a**. **d**, Output of cell type classifier (logistic regression; Methods, *Identification of clusters’ cell types* section). Heatmap showing the fraction of cells classified to known hippocampal cell types (X axis) out of each cell cluster (Y axis). Cluster annotations are as in **a**. Abbreviations: CA1–3, *cornu Ammonis* regions 1–3; DG, dentate gyrus; Ex, excitatory; ExN, excitatory neurons; ExN IEGs, recently activated excitatory neurons expressing immediate early genes (IEGs); GABA, GABAergic neurons; OPCs, oligodendrocyte precursor cells; Sub, subiculum. **e**, **f**, Cell type annotations post filtering. **e**, Dot plot featuring the expression of marker genes (color scale) across cell types and the percentage of cells expressing them (dot size). **f**, Changes in the frequency of neuronal and non-neuronal cell types across experimental conditions (post filtering). Statistical analyses: two-way ANOVA followed by Fisher’s LSD *post hoc* test. Note: the statistics for ExN DG was performed on log-transformed data. Box plots represent the minimum and maximum values (whiskers), the first and third quartiles (box boundaries), the median (box internal line), and the mean (cross).

**Extended Data Fig. 5:**
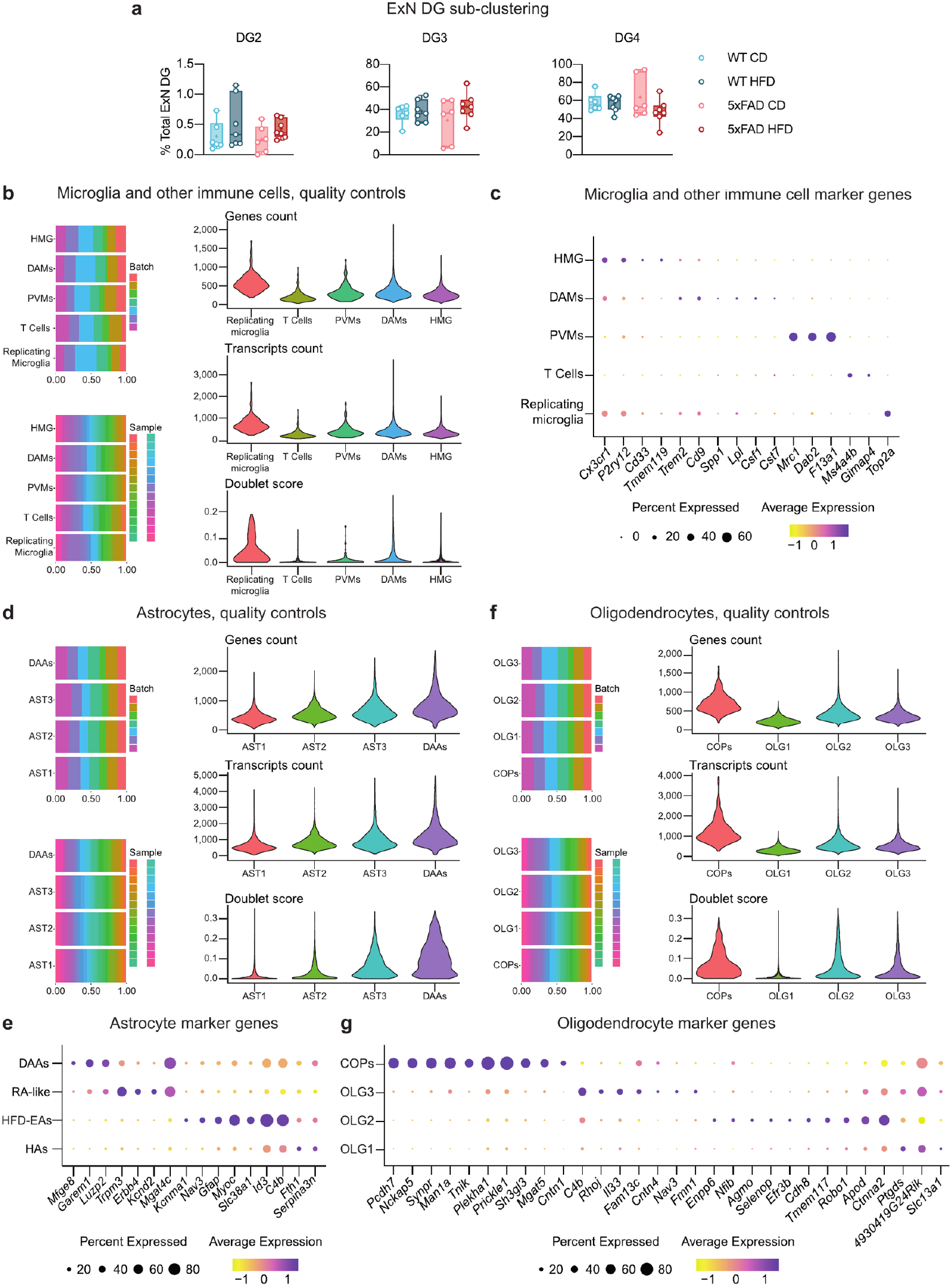
Sub-clustering of DG neurons and glial cells. Mice are the same described in Fig. 1k–m, sample *n*: WT CD=6, WT HFD=7, 5xFAD CD=6, 5xFAD HFD=7. **a**, Changes in frequency of cell states of dentate gyrus granule neurons (ExN DG) across all experimental groups. Sub-clustering analysis identified four clusters (DG1–4), as shown in Fig. 1k; plot relative to DG1 is shown in Fig. 1l. Statistical analyses: two-way ANOVA followed by Fisher’s LSD *post hoc* test; DG4 – statistics on log-transformed data. Box plots represent the minimum and maximum values (whiskers), the first and third quartiles (box boundaries), the median (box internal line), and the mean (cross). **b**, Quality controls for microglia and other immune cells sub-clustering analysis. Abbreviations: DAM, disease-associated microglia; HMG, homeostatic microglia; PVMs, perivascular macrophages. Left: The percent of cells per cluster, from the different batches (top) and from each sample (bottom). Right: violin plots showing the distribution of number of genes (top), number of transcripts (middle), and doublet score (bottom) detected in each cluster. **c**, Dot plot featuring the expression of marker genes (color scale) and the percentage of cells expressing them (dot size) across microglia states and other immune cell types. **d**–**g**, Quality controls for astrocytes (**d**, **e**) and oligodendrocytes (**f**, **g**) sub-clustering analysis. Abbreviations: AST1–3, astrocyte clusters 1–3; COPs, committed oligodendrocyte precursors; DAAs, disease-associated astrocytes; OLG1–3, oligodendrocyte clusters 1–3. Left: percent of cells per cluster, from the different batches (top) and from each sample (bottom). Right: violin plots showing the distribution of number of genes (top), number of transcripts (middle), and doublet score (bottom) detected in each cluster. **e**, **g**, Dot plots featuring the expression of marker genes (color scale) and the percentage of cells expressing them (dot size) of astrocytes (**e**) and oligodendrocytes (**g**).

**Extended Data Fig. 6:**
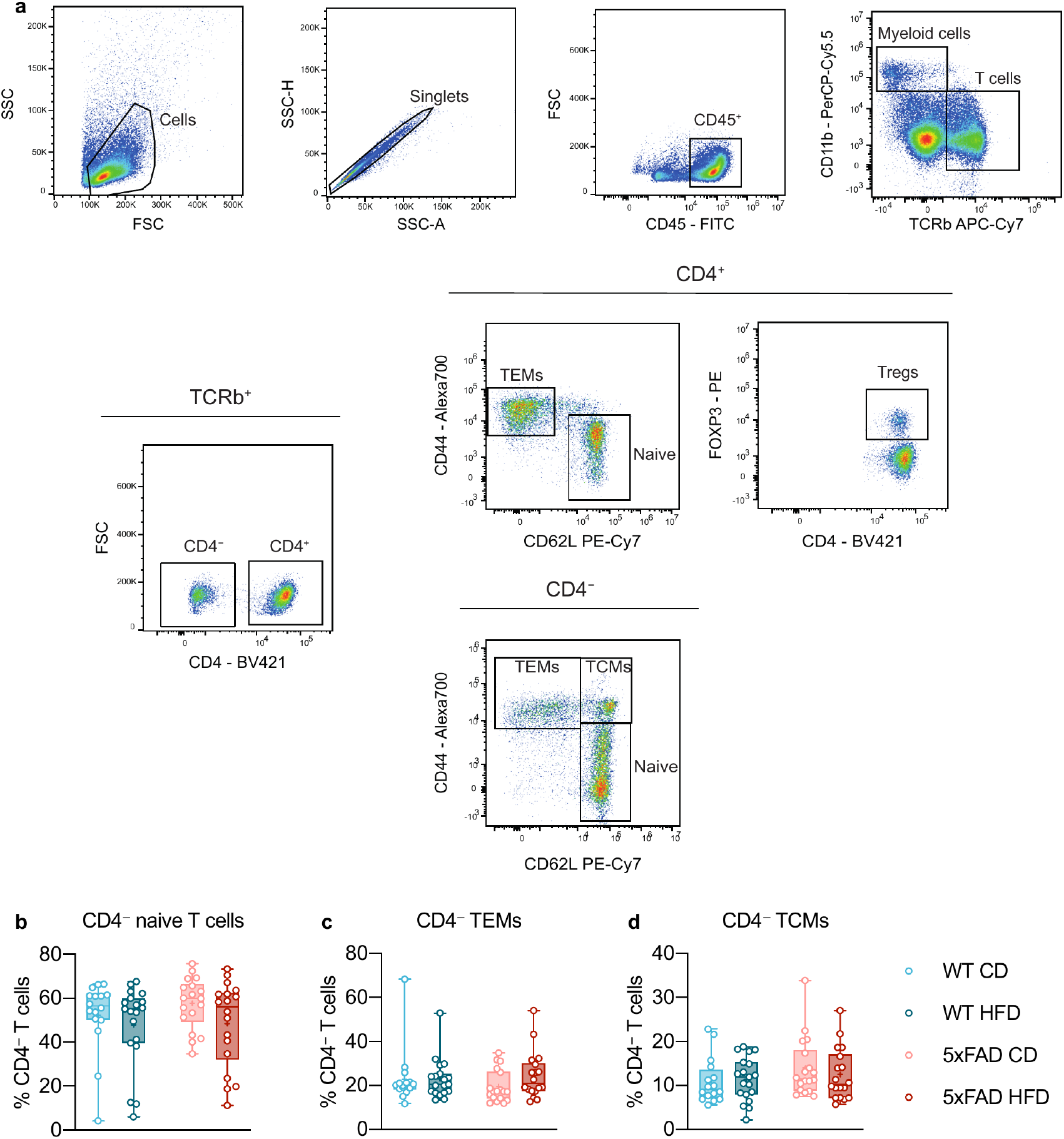
HFD does not affect the systemic CD4^−^ T-cell compartment. Mice are the same described in Fig. 2a–c, sample *n*: WT CD=16, WT HFD=19, 5xFAD CD=18, 5xFAD HFD=18. **a**, Flow cytometric characterization of the CD4^+^ and CD4^−^ T-cell compartments, gating strategy. **b**–**d**, Quantification of splenic frequencies of CD4^−^ naive T cells (CD44^low^CD62L^high^; **b**), CD4^−^ TEMs (CD44^high^CD62L^low^; **c**), and CD4^−^ TCMs (CD44^high^CD62L^high^; **d**). Statistical analyses: two-way ANOVA followed by Fisher’s LSD *post hoc* test. Note: the statistics for CD4^−^ TEMs (**c**) and CD4^−^ TCMs (**d**) was performed on log-transformed data. Box plots represent the minimum and maximum values (whiskers), the first and third quartiles (box boundaries), the median (box internal line), and the mean (cross).

**Extended Data Fig. 7:**
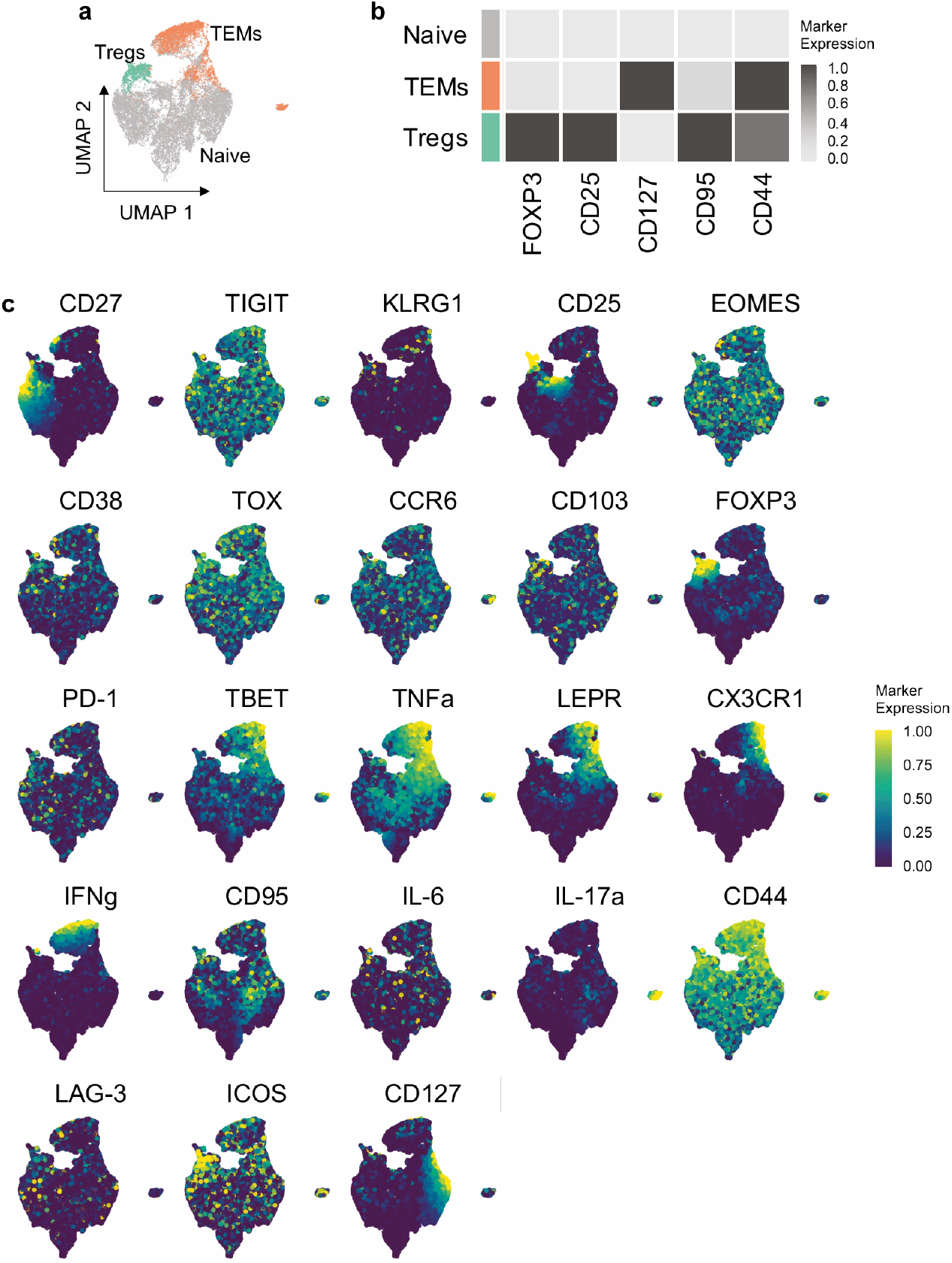
Single-cell profiling of CD4^+^ splenocytes by mass cytometry. Mice are the same described in Fig. 2e–f, sample *n*: WT CD=5, WT HFD=5, 5xFAD CD=5, 5xFAD HFD=5. **a**, UMAP embedding of CD4^+^ cell clusters (2,000 cells, randomly selected from each animal). FlowSOM-based immune populations are overlaid as a color dimension. **b**, Average expression levels of the markers used for UMAP visualization and FlowSOM clustering. **c**, The expression of each indicated marker is overlaid.

**Extended Data Fig. 8.**
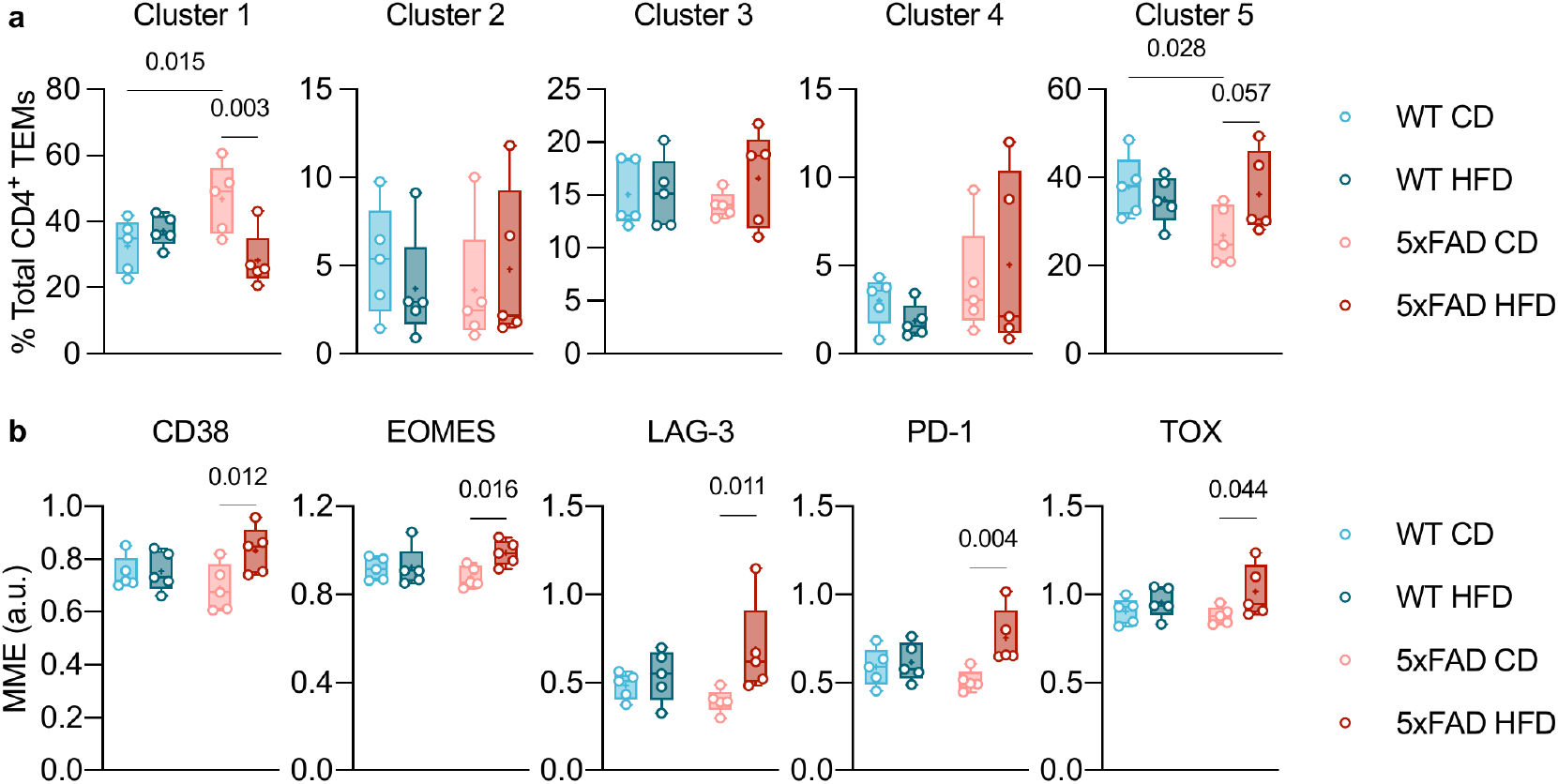
Sub-clustering analysis of the CD4^+^ TEM compartment and expression of selected exhaustion-related markers. Mice are the same described in Fig. 2e–f, sample *n*: WT CD=5, WT HFD=5, 5xFAD CD=5, 5xFAD HFD=5. **a**, Sub-clustering analysis of the CD4^+^ TEM compartment identified six clusters; plot relative to Cluster 6 is reported in Fig. 2f. **b**, HFD increases the expression of exhaustion markers in CD4^+^ TEMs in 5xFAD mice. Expression of selected exhaustion-related markers. MME, mean marker expression. Statistical analyses: two-way ANOVA followed by Fisher’s LSD *post hoc* test. Note: the statistics for Cluster 2 (**a**), Cluster 3 (**a**), Cluster 4 (**a**) and TOX (**b**) was performed on log-transformed data. Box plots represent the minimum and maximum values (whiskers), the first and third quartiles (box boundaries), the median (box internal line), and the mean (cross).

**Extended Data Fig. 9:**
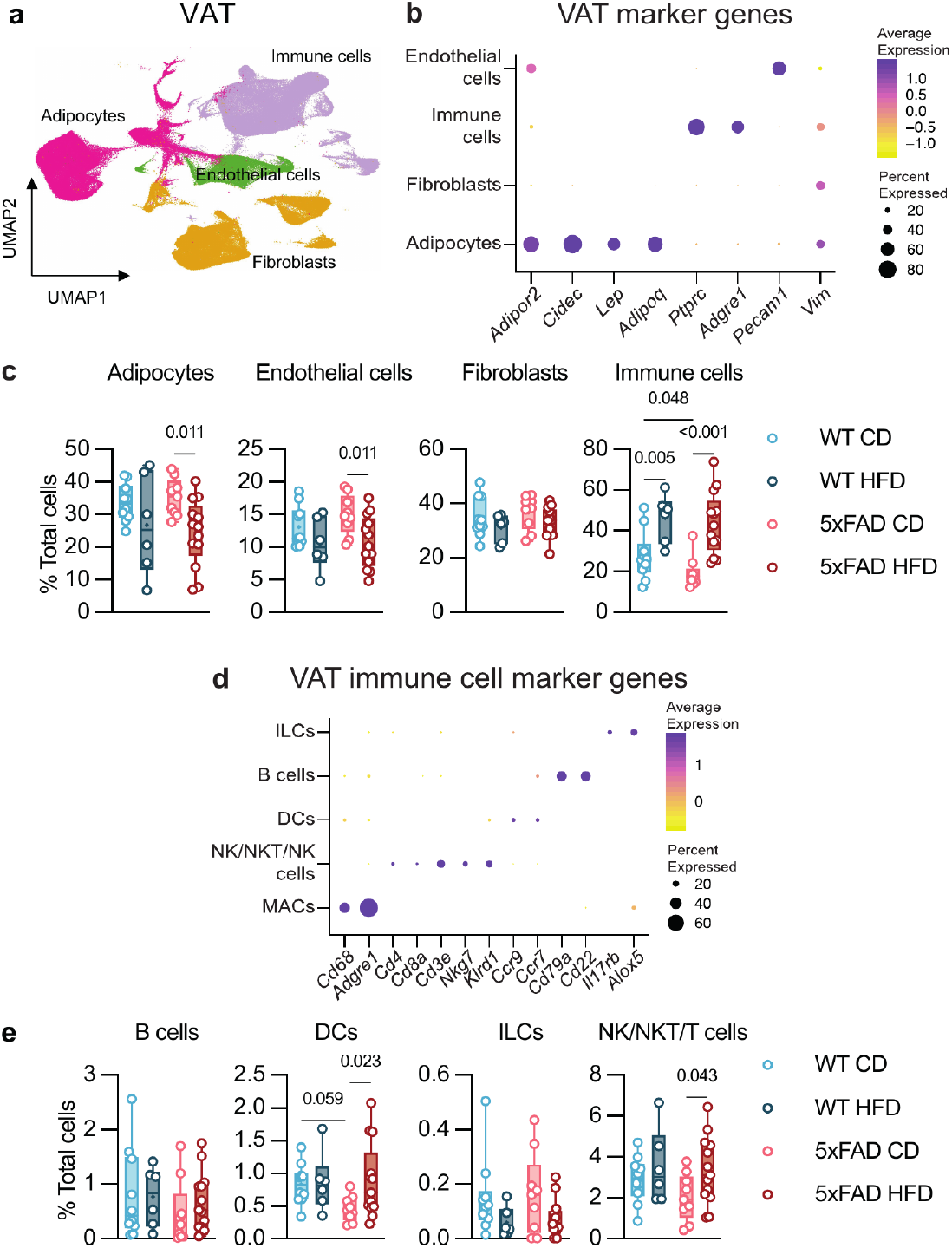
sNuc-Seq of the mouse VAT. Mice are the same described in Fig. 4a–d, sample *n*: WT CD=10, WT HFD=6, 5xFAD CD=9, 5xFAD HFD=13. **a**–**c**, Main cell populations in the mouse gonadal VAT across all genotype and diet conditions. **a**, UMAP embedding of single nucleus profiles (sNuc-Seq), colored after *post hoc* cell type annotation. **b**, Dot plot featuring the expression of marker genes (color scale) and the percentage of cells expressing them (dot size) of VAT cell types. **c**, Changes in frequency of cell types across experimental conditions. **d**, **e**, Sub- clustering analysis of VAT immune cells. **d**, Dot plot featuring the expression of marker genes (color scale) and the percentage of cells expressing them (dot size) of VAT immune cell types. **e**, Changes in frequency of cell types across experimental conditions. Plot relative to MACs is shown in Fig. 4b. Abbreviations: DCs, dendritic cells; ILCs, innate lymphoid cells; MACs, macrophages; NK, natural killer cells; NKT, natural killer T cells. **c**, **e**, Statistical analyses: two-way ANOVA followed by Fisher’s LSD *post hoc* test. Note: the statistics for Immune cells (**c**) and B cells (**e**) was performed on log-transformed data; the statistics for ILCs (**e**) was performed on log(1+x)- transformed data. Box plots represent the minimum and maximum values (whiskers), the first and third quartiles (box boundaries), the median (box internal line), and the mean (cross).

**Extended Data Fig. 10:**
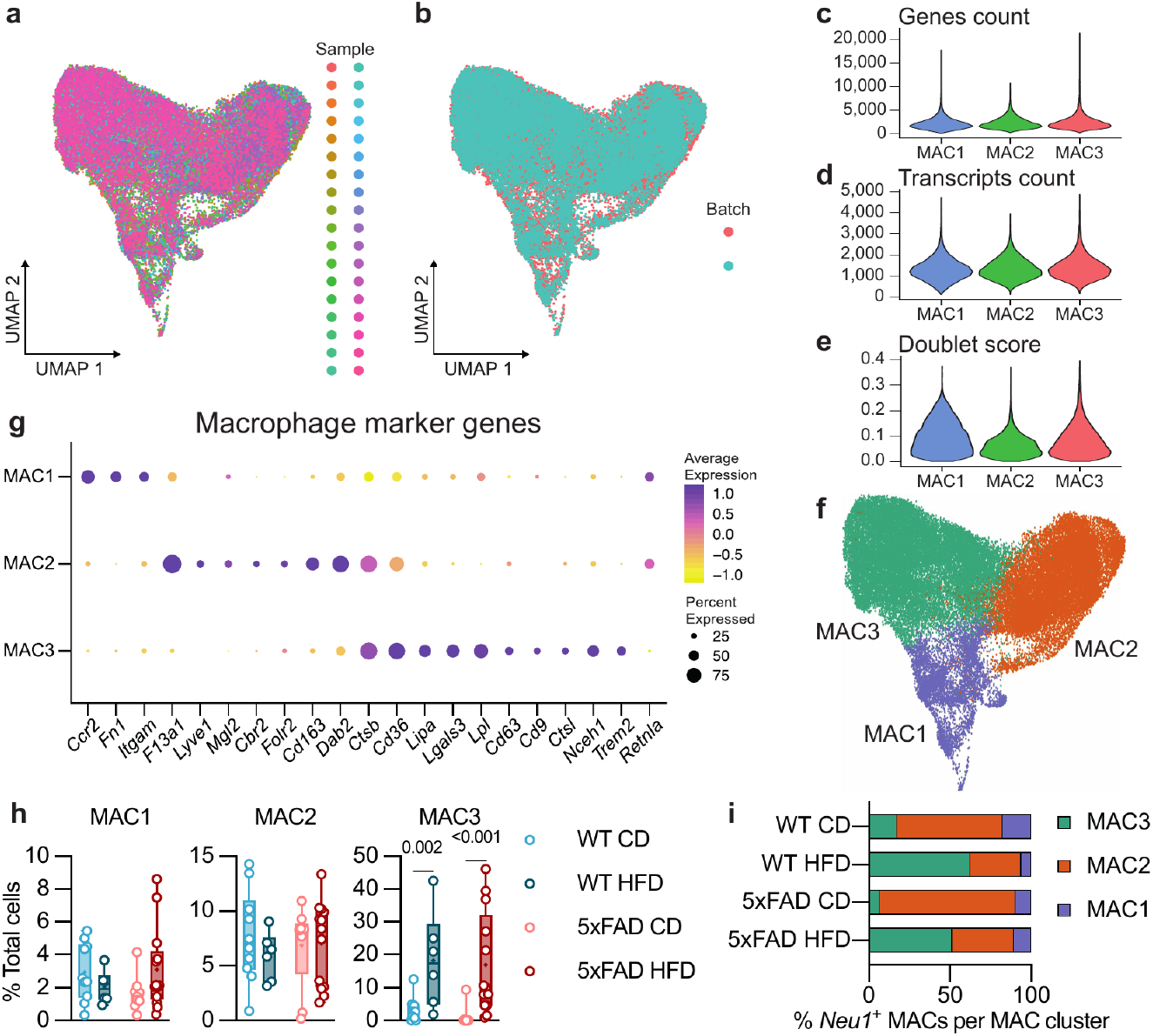
Sub-clustering analysis of VAT macrophages. Mice are the same described in Fig. 4a–d, sample *n*: WT CD=10, WT HFD=6, 5xFAD CD=9, 5xFAD HFD=13. Three MAC clusters were identified, annotated as MAC1–3. **a**–**e**, Quality controls for adipose macrophages sub-clustering analysis. **a**, **b**, Macrophage cells in adipose tissue cluster independently of sample (**a**) and batch effects (**b**). **c**–**e**, Violin plots showing the distribution of number of genes (**c**), number of transcripts (**d**), and doublet scores (**d**) detected in each MAC cluster. **f**, VAT MAC clusters, UMAP embedding of single nucleus profiles (sNuc-Seq), colored after *post hoc* cell type annotation. **g**, Dot plot featuring the expression of marker genes (color scale) and the percentage of cells expressing them (dot size) of VAT MACs. **h**, Changes in frequency of MAC clusters across experimental conditions. Statistical analyses: two-way ANOVA followed by Fisher’s LSD *post hoc* test. Note: the statistics for and MAC1 was performed on log-transformed data; the statistics for MAC3 was performed on log(1+x)-transformed data. Box plots represent the minimum and maximum values (whiskers), the first and third quartiles (box boundaries), the median (box internal line), and the mean (cross). **i**, Stacked bar plot showing the distribution of *Neu1*-expressing macrophages in each macrophage cluster across experimental conditions. Note: in one of the 5xFAD CD mice, *Neu1*-expressing macrophages were below detection level in all MAC clusters.

**Extended Data Fig. 11:**
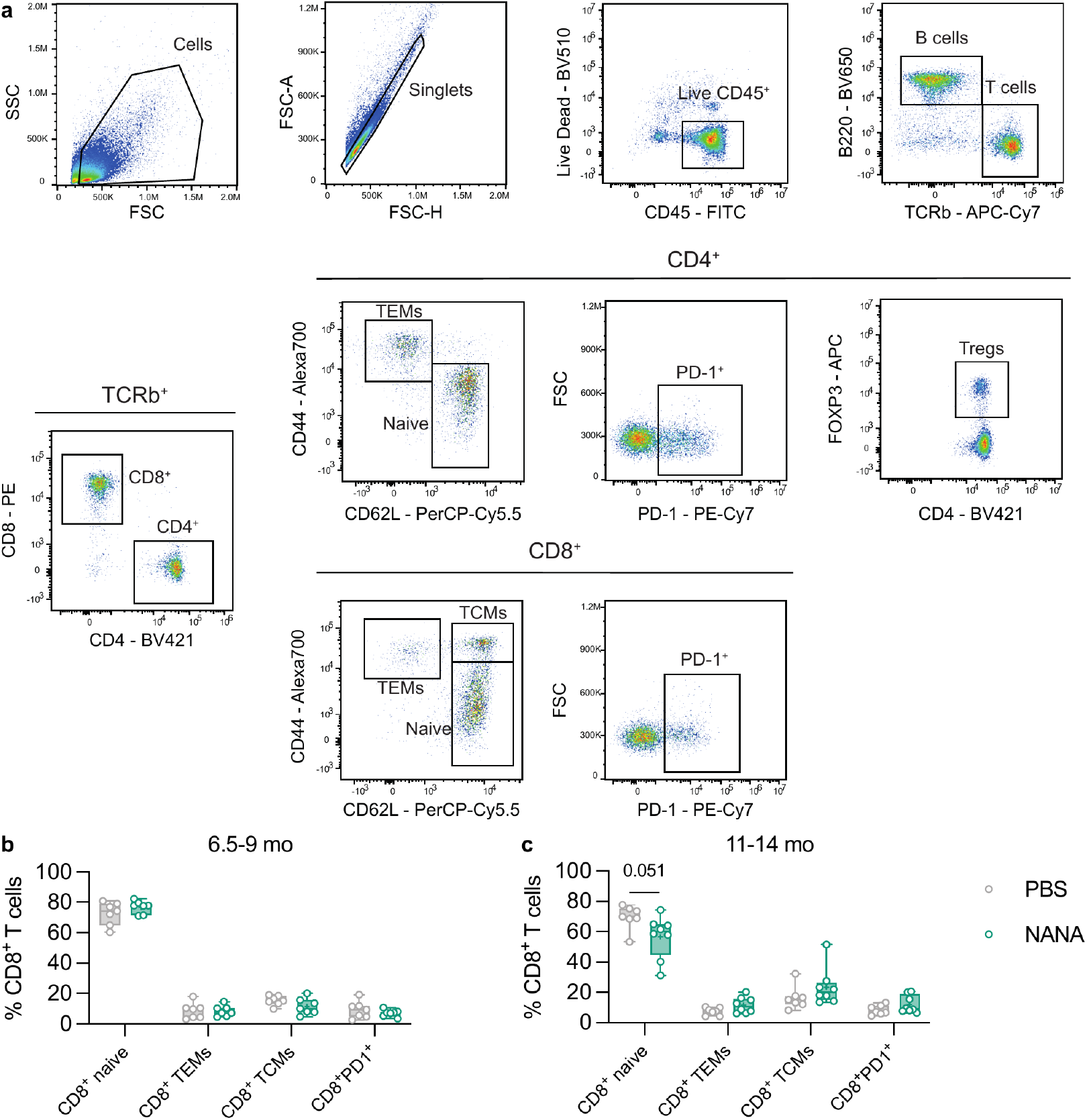
NANA treatment does not affect young adult mice or the CD8^+^ T-cell compartment in elderly mice. **a**, Flow cytometric characterization of the CD4^+^ and CD8^+^ T-cell compartments, gating strategy. **b**, **c**, Quantification of splenic frequencies of CD8^+^ naive T cells (CD44^low^CD62L^high^), CD8^+^ TEMs (CD44^high^CD62L^low^), CD8^+^ TCMs (CD44^high^CD62L^high^), and CD8^+^PD1^+^ T cells in young adult (**b**) and middle-aged mice (**c**). Mice are the same described in Fig. 5b, c, sample *n*: **b**, PBS=7, NANA=7; **c**, PBS=7, NANA=8. Statistical analyses: multiple two-tailed unpaired *t*-tests. Box plots represent the minimum and maximum values (whiskers), the first and third quartiles (box boundaries), the median (box internal line), and the mean (cross).

**Extended Data Fig. 12:**
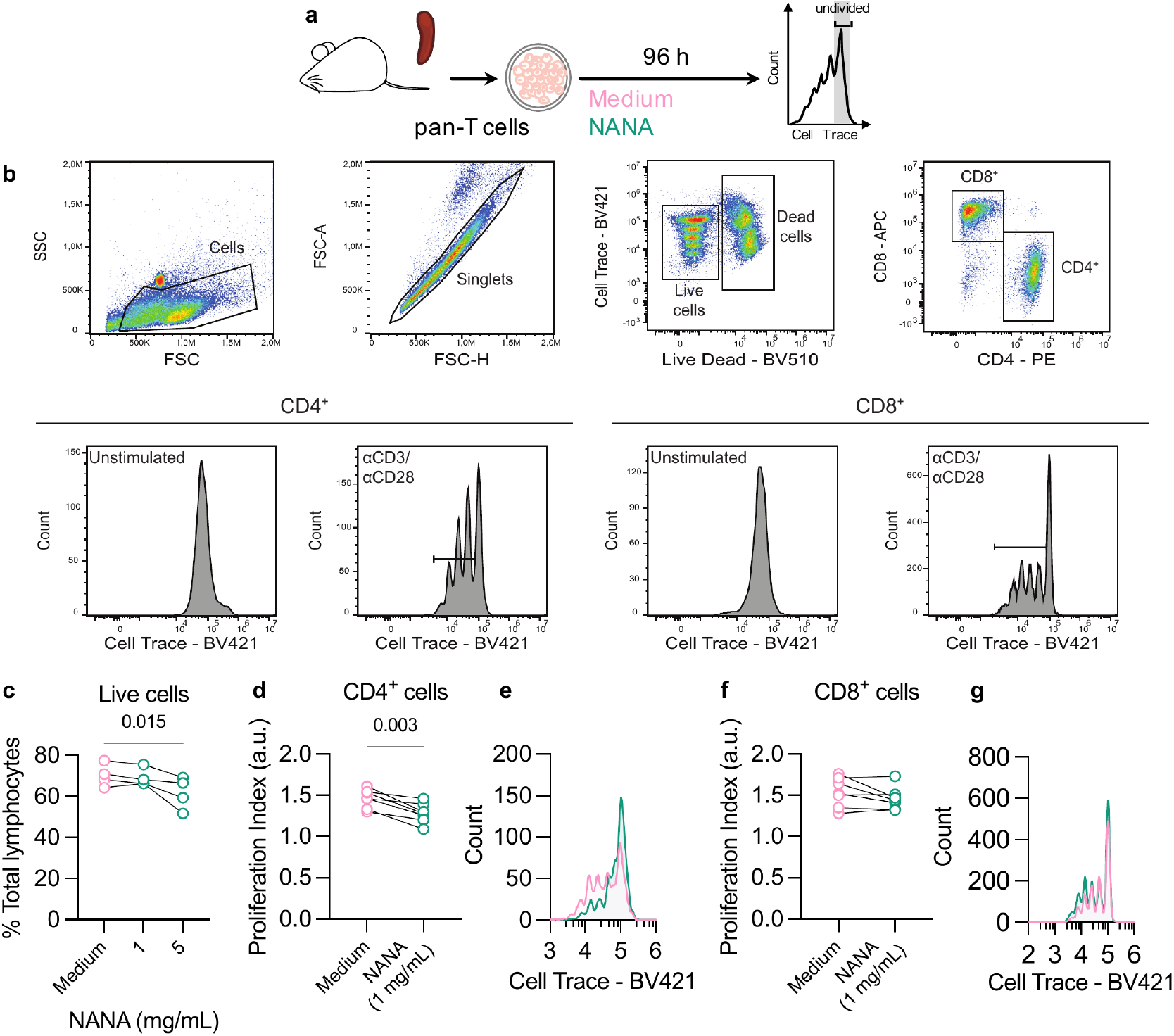
NANA treatment impairs mouse CD4^+^ T-cell proliferation *in vitro*. **a**, Schematic of treatment regimen in mouse pan-T-cell cultures and proliferation assay. **b**, Flow cytometric analysis of CD4^+^ and CD8^+^ T-cell proliferation, gating strategy. **c**, Viability assay with two different doses of NANA. Data from a single experiment, sample *n*=4 mice, three conditions (NANA 1 mg/mL, NANA 5 mg/mL, and medium control) for each. Each black line represents one mouse and connects the three tested conditions. Statistical analyses: one-way within-subjects ANOVA followed by Fisher’s LSD *post hoc* test. **d**, NANA exposure reduces proliferation of CD4^+^, but not CD8^+^ T cells. Assessment of mouse CD4^+^ T-cell proliferation ability. **e**, Histograms of a representative sample. **f**, Assessment of mouse CD8^+^ T-cell proliferation. **g**, Histograms of a representative sample. **d**–**g**, Data from two independent experiments, of which one is the same described in **c**, sample *n*=7 mice, two conditions (NANA 1 mg/mL and medium control) for each. Each black line represents one mouse and connects the two tested conditions. Statistical analyses: paired two-tailed *t*-test.

**Extended Data Fig. 13:**
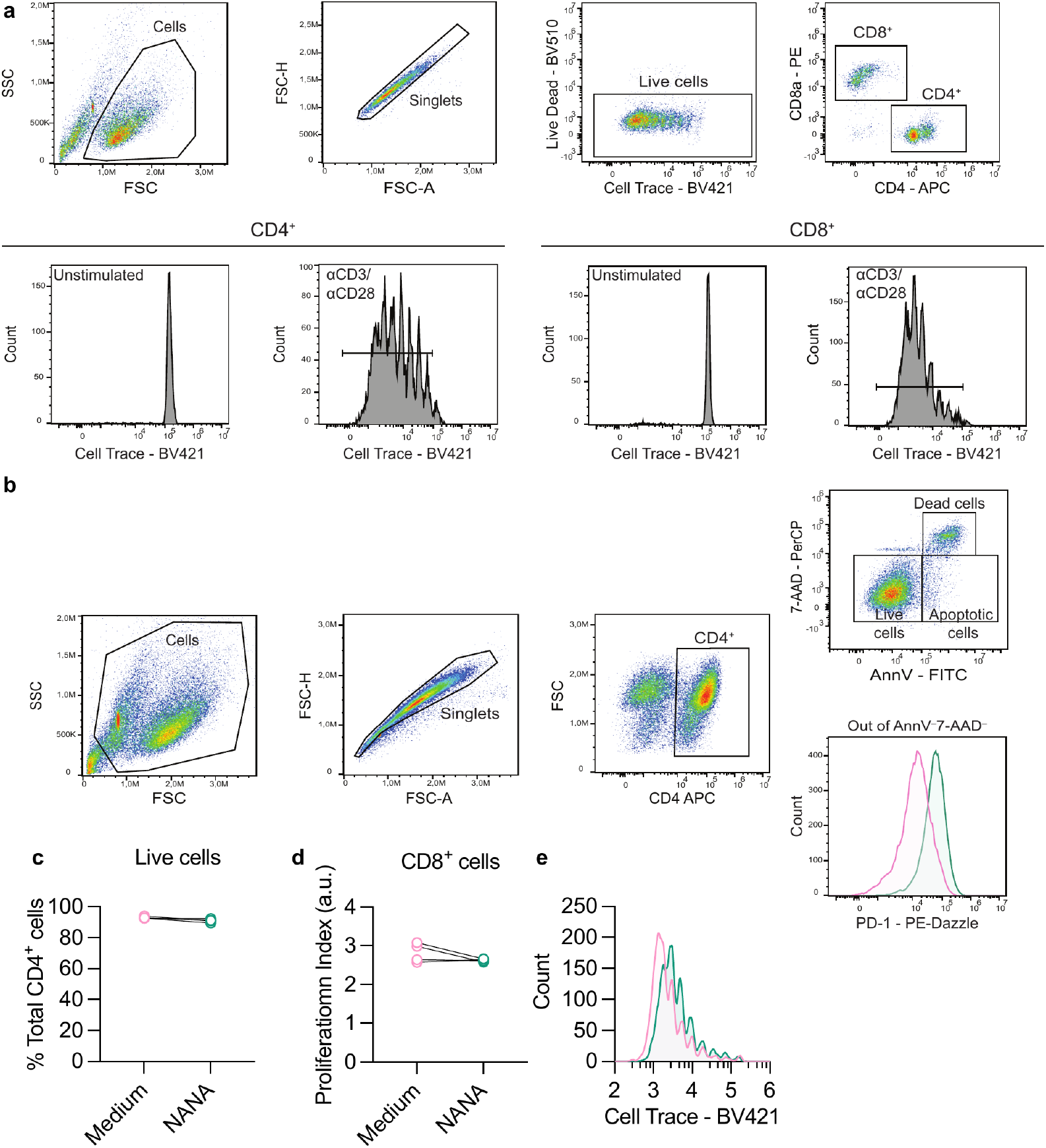
NANA treatment has no impact on human CD4^+^ T-cell viability and CD8^+^ T-cell proliferation. **a**, **b**, Flow cytometric analysis of human CD4^+^ and CD8^+^ T-cell proliferation (**a**) and apoptosis/viability assay (**b**), gating strategy. **b**, PD-1 expression (geometric mean fluorescent intensity) was evaluated in annexin V (AnnV)/7-amino-actinomycin D (7- AAD)-double negative cells. **c**, NANA exposure does not affect CD4^+^ T-cell viability. **d**, NANA exposure does not affect human CD8^+^ T-cell proliferation. **e**, Histograms of a representative sample. **c**, **d**, Data refer to the analysis of the same samples described in Fig. 5d–h, sample *n*=4 human individuals (blood). Each black line represents one individual and connects the two tested conditions. Statistical analyses: paired two-tailed *t*-test.

**Extended Data Fig. 14:**
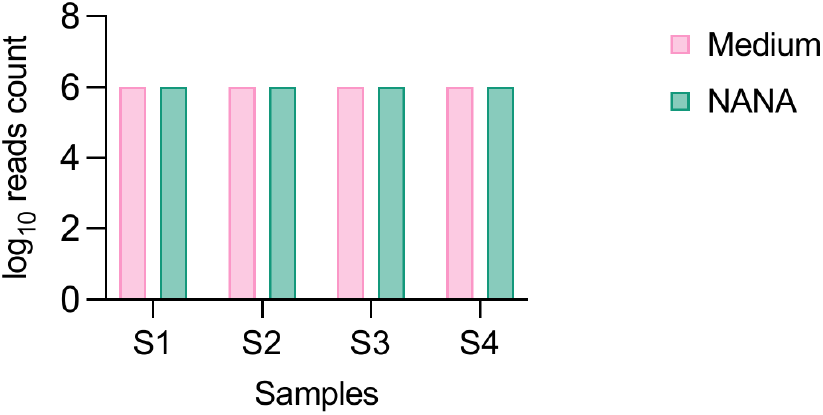
Quality control for bulk RNA-Seq of human T-cell cultures. Reads count per sample (log_10_). For each sample, 24,594 genes were used. Data refer to the analysis of the same samples described in Fig. 5d–h, sample *n*=4 human individuals (blood). S1–4: subject 1– 4.

**Extended Data Table 1.** Gene set enrichment analysis of the dentate gyrus cluster DG1 after sNuc-Seq of the mouse hippocampus.

**Extended Data Table 2.** Differentially regulated genes (DESeq2, FDR <0.050) in human T-cell cultures after NANA treatment relative to medium-treated controls.

**Extended Data Table 3.** Gene set enrichment analysis (FDR <0.050) of human T-cell cultures after NANA treatment relative to medium-treated controls.

**Extended Data Table 4.** Fluorochrome-labelled monochrome antibodies used in the study.

| Antigen | Fluorophore | Dilution | Supplier | Cat. # |
| --- | --- | --- | --- | --- |
| CD62L | PE-Cy7 | 1:150 | Biolegend | 269976 |
| CD11b | PerCP-Cy5.5 | 1:200 | Biolegend | 276002 |
| CD44 | AlexaFluor700 | 1:150 | Biolegend | 292685 |
| TCRb | APC-Cy7 | 1:150 | Biolegend | 281644 |
| CD4 | BV421 | 1:150 | Biolegend | 288315 |
| CD45 | FITC | 1:150 | Biolegend | 228450 |
| FOXP3 | PE | 1:150 | Biolegend | 271499 |
| B220 | BV650 | 1:150 | Biolegend | 308321 |
| CD8a | PE | 1:150 | Biolegend | 100708 |
| PD-1 | PE-Cy7 | 1:100 | Biolegend | 266040 |
| FOXP3 | APC | 1:100 | Invitrogen | 17-5773-82 |
| CD62L | PerCP-Cy5.5 | 1:150 | Biolegend | 289783 |
| CD4 | APC | 1:150 | Biolegend | 166838 |
| CD4 | PE | 1:150 | Biolegend | 196678 |
| CD8a | APC | 1:200 | Biolegend | 288881 |
| CD8a | PE | 1:200 | Biolegend | 250825 |
| CD4 | APC | 1:200 | Biolegend | 267985 |
| PD-1 | PE-Dazzle | 1:100 | Biolegend | 264152 |

**Extended Data Table 5.** Heavy-metal-conjugated antibodies used in the study.

| Antigen | Metal | Supplier | Cat. # | In-house conjugation | Staining step |
| --- | --- | --- | --- | --- | --- |
| CD45 | <sup>89</sup>Y | Fluidigm | 3089005B |  | S |
| CD27 | <sup>113</sup>In | Biolegend | 124202 | yes | S |
| TCRgd | <sup>115</sup>In | R&D | MAB7297 | yes | S |
| TCR-b | <sup>143</sup>Nd | Fluidigm | 3143010B |  | S |
| CCR2 | <sup>144</sup>Nd | R&D | MAB55381R | yes | S |
| CD4 | <sup>145</sup>Nd | Fluidigm | 3145002B |  | S, I |
| TIGIT | <sup>146</sup>Nd | Biolegend | 142102 | yes | S |
| KLRG1 | <sup>147</sup>Sm | R&D | MAB69441 | yes | S |
| CD11b | <sup>148</sup>Nd | Fluidigm | 3148003B |  | S |
| CD19 | <sup>149</sup>Sm | Fluidigm | 3149002B |  | S |
| CD25 | <sup>150</sup>Nd | Fluidigm | 3150002B |  | S |
| CD8 | <sup>153</sup>Eu | Fluidigm | 3153012B |  | S |
| CD38 | <sup>154</sup>Sm | Biolegend | 102702 | yes | S |
| CCR6 | <sup>156</sup>Gd | Fluidigm | 3156016A |  | S |
| CD103 | <sup>157</sup>Gd | Biolegend | 121402 | yes | S |
| PD-1 | <sup>159</sup>Tb | Fluidigm | 3159006B |  | S |
| LEPR | <sup>163</sup>Dy | R&D | AF497 | yes | S |
| CX3CR1 | <sup>164</sup>Dy | Fluidigm | 3164023B |  | S |
| CD95 | <sup>166</sup>Er | R&D | AF435 | yes | S |
| CD44 | <sup>171</sup>Yb | Fluidigm | 3171003B |  | S |
| LAG-3 | <sup>174</sup>Yb | Fluidigm | 3174019B |  | S |
| CD127 | <sup>175</sup>Lu | Fluidigm | 3174013B |  | S |
| ICOS | <sup>176</sup>Yb | Fluidigm | 3176014B |  | S |
| EOMES | <sup>152</sup>Sm | Biolegend | MAB8889 | yes | I |
| TOX | <sup>155</sup>Gd | Thermo Fisher | 14-6502-95 | yes | I |
| FOXP3 | <sup>158</sup>Gd | Fluidigm | 3158003A |  | I |
| TBET | <sup>161</sup>Dy | Fluidigm | 3161014B |  | I |
| TNFα | <sup>162</sup>Dy | Fluidigm | 3162002B |  | I |
| IFNγ | <sup>165</sup>Ho | Fluidigm | 3165003B |  | I |
| IL-6 | <sup>167</sup>Er | Fluidigm | 3167003B |  | I |
| IL-17a | <sup>169</sup>Tm | Fluidigm | 3169005B |  | I |

Abbreviations: S, surface staining; I, intracellular staining.

**Extended Data Table 6.** Raw data, statistical analyses, and graphs utilized in main Figure and Extended Data Figure panels (file generated with GraphPad Prism).

